# Structure of an LGR dimer – an evolutionary predecessor of glycoprotein hormone receptors

**DOI:** 10.1101/2024.12.31.630923

**Authors:** Zhen Gong, Shuobing Chen, Ziao Fu, Brian Kloss, Chi Wang, Oliver B. Clarke, Qing R. Fan, Wayne A. Hendrickson

**Affiliations:** Department of Biochemistry and Molecular Biophysics, Columbia University, New York, NY 10032; New York Structural Biology Center, New York, NY 10027; Department of Physiology and Cellular Biophysics, Columbia University, New York, NY 10032; Department of Anesthesiology, Columbia University Irving Medical Center, New York, NY 10032; Irving Institute for Clinical and Translational Research, Columbia University, New York, NY 10032; Department of Molecular Pharmacology and Therapeutics, Columbia University, New York, NY 10032; Department of Pathology and Cell Biology, Columbia University, New York, NY 10032

## Abstract

The glycoprotein hormones of humans, produced in the pituitary and acting through receptors in the gonads to support reproduction and in the thyroid gland for metabolism, have co-evolved from invertebrate counterparts^1,2^. These hormones are heterodimeric cystine-knot proteins; and their receptors bind the cognate hormone at an extracellular domain and transmit the signal of this binding through a transmembrane domain that interacts with a heterotrimeric G protein. Structures determined for the human receptors as isolated for cryogenic electron microscopy (cryo-EM) are all monomeric^3–6^ despite compelling evidence for their functioning as dimers^7–10^. Here we describe the cryo-EM structure of the homologous receptor from a neuroendocrine pathway that promotes growth in a nematode^11^. This structure is an asymmetric dimer that can be activated by the hormone from that worm^12^, and it shares features especially like those of the thyroid stimulating hormone receptor (TSHR). When studied in the context of the human homologs, this dimer provides a structural explanation for the transactivation evident from functional complementation of binding-deficient and signaling-deficient receptors^7^, for the negative cooperativity in hormone action that is manifest in the 1:2 asymmetry of primary TSH:TSHR complexes^8,9^, and for switches in G-protein usage that occur as 2:2 complexes form^9,10^.

## Introduction

There are three glycoprotein hormone receptors (GpHRs) in humans: luteinizing hormone– choriogonadotropin receptor (LHCGR), follicle stimulating hormone receptor (FSHR) and TSHR. Together, they play important roles in reproduction and regulating metabolism^1^. GpHRs belong to a subfamily of class A G protein coupled receptors (GPCRs) called leucine-rich-repeat-containing GPCRs (LGRs). An LGR is characterized by a large extracellular domain (ECD) with multiple leucine-rich repeats (LRRs), a rhodopsin-like transmembrane domain (TMD) of seven transmembrane (TM) helices, and a disulfide-bridged hinge segment that connects the two. The ECD binds the hormone^13,14^ while the TMD transmits the signal of this binding to cytoplasmic effectors, notably G proteins^15^.

The structures of signaling complexes for all three human GpHRs have been solved recently using cryogenic electron microscopy (cryo-EM)^3–6^. The three hormone-receptor-G-protein complexes are similar to one another; however, in each case the receptor is monomeric, whereas compelling evidence implicates receptor dimers (possibly other oligomers) in GpHR signaling. Förster and bioluminescence resonance energy transfer (FRET and BRET)^8,16–18^ and super-resolution microscopy experiments^19^ demonstrate associations; negative cooperativity in hormone binding as well as assays of binding competition for chimeric receptors imply asymmetric 1:2 hormone:receptor association^8,10,20^; switches in TSHR signaling from being through Gs to produce cAMP to being through Gq to produce inositol phosphate^10^ or through Gi/o for a biphasic cAMP response, which in consort require the formation of 2:2 TSH:TSHR complexes^9,21^; and functional complementation of mutants defective in binding by mutants defective in signaling require transactivation from one protomer to another^8,22–24^. Most impressively, transgenic mice co-expressing binding-deficient and signaling-deficient LHCGRs reinstate functional hormone action in the absence of wild-type receptors^7^. Such complementation is intrinsic to the natural functioning of the type C heterodimeric GABA_B_ receptors^25,26^.

The glycoprotein hormones and their receptors co-evolved from homologs in invertebrate metazoans^27^, in which the hormones are α2/β5 heterodimers and their type A LGR receptors are already distinguished as having 12 LRRs bounded by disulfide-bridged regions at the N-terminus and at the hinge region^28^. The *Caenorhabditis elegans* genome includes a single LGR gene^29^ and genes for the α2^30^ and β5^31^ chains encode the *Ce*LGR ligand (*Ce*α2β5); gene knockouts of any one of these produce the same growth defective phenotype in the nematode^11^. The human homolog of the *Ce*LGR ligand can activate human TSHR, and it is called thyrostimulin (*Hs*α2β5)^32^.

We previously solved the crystal structure of *Ce*α2β5^12^, which has expected similarity to human glycoprotein hormones but is unglycosylated. In this study, we report a nearly full-length cryo-EM structure of the *C. elegans* LGR, which is an asymmetric dimer in an unliganded state. We also show that *Ce*α2β5 is able to activate *Ce*LGR to produce cyclic adenosine monophosphate (cAMP) in human embryonic kidney (HEK) cells.

## Results

### Characterization of the *Ce*LGR receptor

The full-length *Ce*LGR contains 929 residues, including a C-terminal tail extending beyond the limits of hGpHRs and predicted by AlphaFold to be unstructured (sequence alignment in Extended Data Fig. 1). For protein production, we truncated residues Ile713 to Ser929 of *Ce*LGR after helix 8 and replaced its native signal peptide (residues 1 to 30) with an influenza hemagglutinin (HA) signal peptide to improve the expression of *Ce*LGR in HEK293S cells. The purified *Ce*LGR protein without reducing agent appears as monomer and ‘dimer’ bands on an SDS-PAGE gel. In the presence of a reducing agent, the dimer band almost disappears, residual monomer bands persist, and two new bands corresponding to lower molecular weight appear (Extended Data Fig. 2a). Complete cleavage into two subunits was observed for TSHR from human thyroid membranes^33^; but we observe an incomplete cleavage of hTSHR as produced from HEK293S cells, which can be completed by limited trypsin digestion (Extended Data Fig. 2b). Based on evidence that a double cleavage had excised a segment of the hinge region^34^, both cryo-EM structures of hTSHR employed constructs that deleted residues Ala317-Phe366^4,5^.

We used proteolytic mass spectrometry to delimit the cleavage site that generated the two polypeptides found in our preparations of *Ce*LGR and of hTSHR. We analyzed each protein after reduction and subsequent digestion with three proteases, coupling the observed coverages from identified proteolytic peptides with knowledge of contiguously ordered cryo-EM structure on either side of a disordered segment, which is where the natural cleavages must have occurred. These considerations delimit the cleavage of *Ce*LGR to within residues Arg343-Arg354 and that of hTSHR to within its residues Arg310-Ser314 (Extended Data Fig. 3). This is surprising for *Ce*LGR since cleavage would have been expected after Arg354 at a canonical furin site (RRKR↓) and surprising for hTSHR in that it excludes the previously suggested cleavages after Asn316 and Phe366^34^. These severances cut at similar sites between the disulfide-bridged cysteine residues of the hinge-region, just beyond HH2 and near the C-terminal end of the hormone-binding ECD (Extended Data Fig. 1). The highly conserved P10 segment joins the ECD to the TMD.

### Cryo-EM structure of the unliganded *Ce*LGR dimer

The structure of *Ce*LGR was determined by single-particle cryo-EM to an overall resolution of 3.79 Å (Fig. 1 and Extended Data Fig. 4-6). The cryo-EM data collection, refinement and validation statistics are summarized in Extended Data Table 1. The structure of *Ce*LGR is an asymmetric homodimer. By comparison to the known structures of hGpHRs^3–6^, the ECDs of both *Ce*LGR protomers are tilted down towards the cell membrane, occupying an inactive conformation. Overall, the two protomers are similar in structure (Extended Data Fig. 7a); but protomer B embraces protomer A after an anti-clockwise rotation of 142° and a translation of 4.1 Å toward the ECD along the rotational screw axis that penetrates the dimeric interface (Fig. 1b and c). The angle between the rotation axis and the normal to cell membrane is 8.3° (Fig. 1b) based on a membrane boundary calculated using the Positioning of Proteins in Membrane (PPM 3.0) server^35^. The two protomers are not identical; rather, ECD and TMD are rigid bodies linked such that the ECD is rotated 7.3° further away from the TMD in protomer A than in B (Extended Data Fig. 7).

**Fig. 1.**
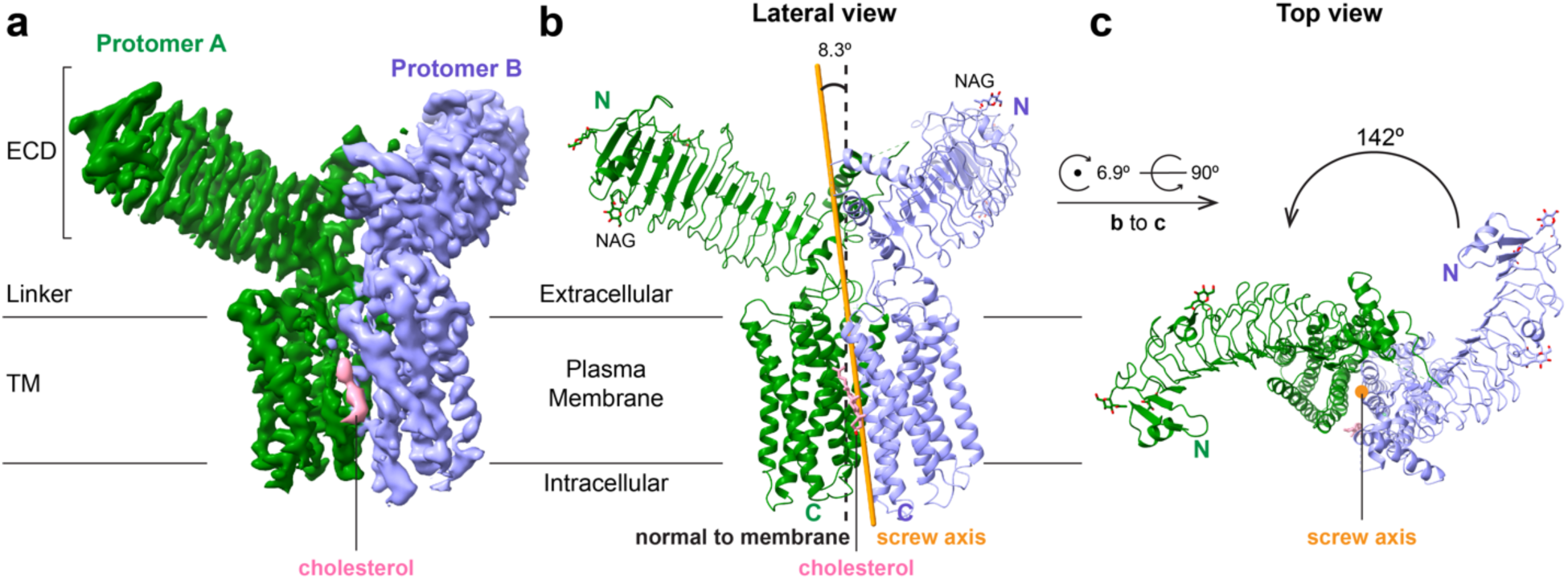
Cryo-EM structure of the *Ce*LGR homodimer in an unliganded apo state. **(a)** Cryo-EM density map of the *Ce*LGR homodimer at an overall resolution of 3.79 Å. **(b)** Lateral view of a ribbon diagram of the atomic model fitted to the cryo-EM density for the *Ce*LGR structure (same view as in **a**). The orange stick is the screw axis relating protomer B to protomer A. *N*-acetylglucosamines (NAGs) and cholesterol are shown as sticks. **(c)** Top view of the *Ce*LGR structure in ribbon diagram, looking directly down along the screw axis (orange stick). Coloring has protomer A (green), protomer B (light purple) and assigned cholesterol (pink).

The density map is well resolved for both protomers from the N termini through TM5. Side chains of residues in these regions could be assigned with certainty. However, despite extensive attempts at classification to resolve ambiguities, the map remains poor at TM6, TM7 and helix 8 for both protomers with little side chain definition. This may be due to a dynamic character for the TM6-TM7 interface. Therefore, the model for TM6 and TM7 in both protomers was built by rigid-body fittings of the AlphaFold-predicted model into the density map.

Five potential N-linked glycosylation sites (Asn80, Asn92, Asn155, Asn372 and Asn410) in *Ce*LGR are predicted by the NetNGlyc-1.0 server^36^. Density was observed to model carbohydrate residues at three potential glycosylation sites (Asn80, Asn92 and Asn155); Asn 372 is in a disordered segment, and density observed at Asn 410 in protomer A was too weak to model.

Since the overall structures of the two protomers are similar and protomer A has the better density, we use it to illustrate structural features in the protomer.

### The N-terminal cysteine-rich and C-terminal cysteine-rich hinge regions

As for the hGpHRs, the extracellular ligand-binding domain of *Ce*LGR consists of 12 LRRs flanked by N-terminal and C-terminal cysteine-rich regions (NCR and CCR, respectively). Disulfide-bridged flanking regions are common features of LRR proteins, thought to serve LRR integrity by capping its ends, and those of *Ce*LGR conform to cysteine motifs that characterize GpHRs^37^. LRRs belong to an archaic prokaryotic protein architecture that is widely used in protein-protein interactions. In eukaryotes, LRR domains developed into key recognition modules in innate immunity^38^.

As for the hGpHRs, the NCR region of *Ce*LGR has an additional β strand, named β0, arranged antiparallel to the LRR β strands that form the inner concave binding surface (Fig. 2a). The NCR region of *Ce*LGR includes three disulfide bridges (Fig. 2a and 2c). The β strand in LRR1 includes three consecutive cysteine residues (Cys67, Cys68 and Cys69), which connect to β0 through the disulfide bonds of Cys69:Cys54 (as for hGpHRs) and Cys67:Cys56; and to the unique N-terminal helix α0 through the disulfide bond of Cys68:Cys43 (Fig. 2a and 2c).

**Fig. 2.**
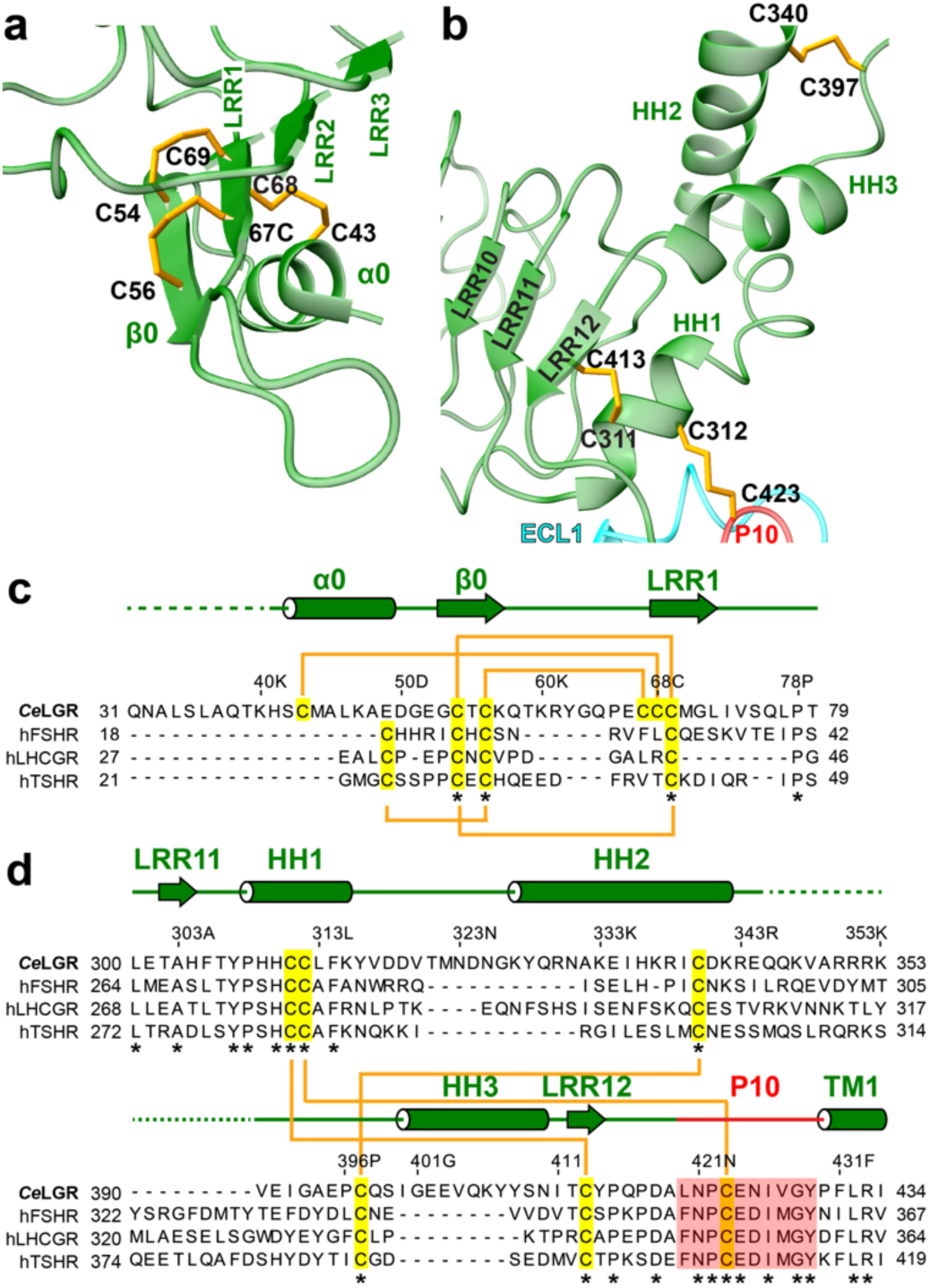
Structure and sequence alignment of the N-terminal cysteine-rich region (a, c) and the C-terminal cysteine-rich hinge region (b, d) of *Ce*LGR protomer A. For the ribbon diagram in **a** and **b**, *N*-acetylglucosamines (NAGs) and *Ce*LGR protomer B are not displayed for clarity. Structural elements and cysteine residues are labeled. The P10 region is colored in red. For the sequence alignment in **c** and **d**, the secondary structural elements are indicated by cylinders for α helices and arrows for β strands. Cysteine residues are highlighted with yellow background, and disulfide bridges are indicated by orange connections. The conserved residues are underscored by asterisks (*). Regions that are unstructured in the model are indicated with green dash lines.

As first described by Jiang et al. for FSHR^14^ and recently confirmed with structures of full-length hGpHRs^3–6^, the CCR hinge region is an integral component of LGR ECDs. Mutagenesis and chimeric receptor studies show importance of the CCR hinge region in GpHR function^39^, seeming to act as a fulcrum^40^ in transmitting the signal from hormone binding to the ECD to G-protein activation by the TMD. In essence, the CCR of *Ce*LGR arises as an elaborated return loop between the β strands of LRR11 and LRR12. For *Ce*LGR, the CCR hinge region includes three hinge helices (HH1, HH2 and HH3), LRR12 and the highly conserved 10-residue fragment (P10, residues Leu420 to Tyr429) (Fig. 2b and 2d); this hinge region is disordered between Arg343 and Ile392. It is noteworthy that HH1, P10 and the six cysteine residues in the CCR hinge region are highly conserved in hGpHRs (Fig. 2b and 2d), indicating a common and important biological function of this region. The HH1 of *Ce*LGR links with LRR12 and P10 through two conserved disulfide bridges Cys311:Cys413 and Cys312:Cys423, while HH2 and HH3 of *Ce*LGR are linked by disulfide bridge Cys340:Cys397 (Fig. 2b and 2d). All three disulfides bridge the cleaved segments.

### Interactions between the ECD and TMD domains of *Ce*LGR

The TMD of each *Ce*LGR protomer arranges itself in the seven-transmembrane-helix conformation of canonical GPCRs, with ECL2 covering the extracellular end in this case (Fig. 3a). P10 in the CCR hinge region of *Ce*LGR directly connects the ECD to the TMD, and this covalent linkage is buttressed by interactions with HH1, ECL1, ECL2 and TM2. The backbone carbonyl group of Leu420^P^^10^ forms hydrogen bonds with the backbone NH and the side chain hydroxyl group of Ser582^ECL^^2^ (Fig. 3b). The side chain of Asn421^P^^10^ forms a hydrogen bond with the side chain of Gln505^ECL^^1^ (Fig. 3b). As mentioned previously, P10 interacts with HH1 through a disulfide bridge Cys423^P^^10^:Cys312^HH^^1^ (Fig. 3b). The backbone carbonyl group of Glu424^P^^10^ forms a hydrogen bond with the backbone NH of Asp495^ECL^^1^ (Fig. 3b). In addition, the backbone NHs of Ile426^P^^10^ and Val427^P^^10^ form hydrogen bonds with the side chain of Asp490^TM^^2^ (Fig. 3b).

**Fig. 3.**
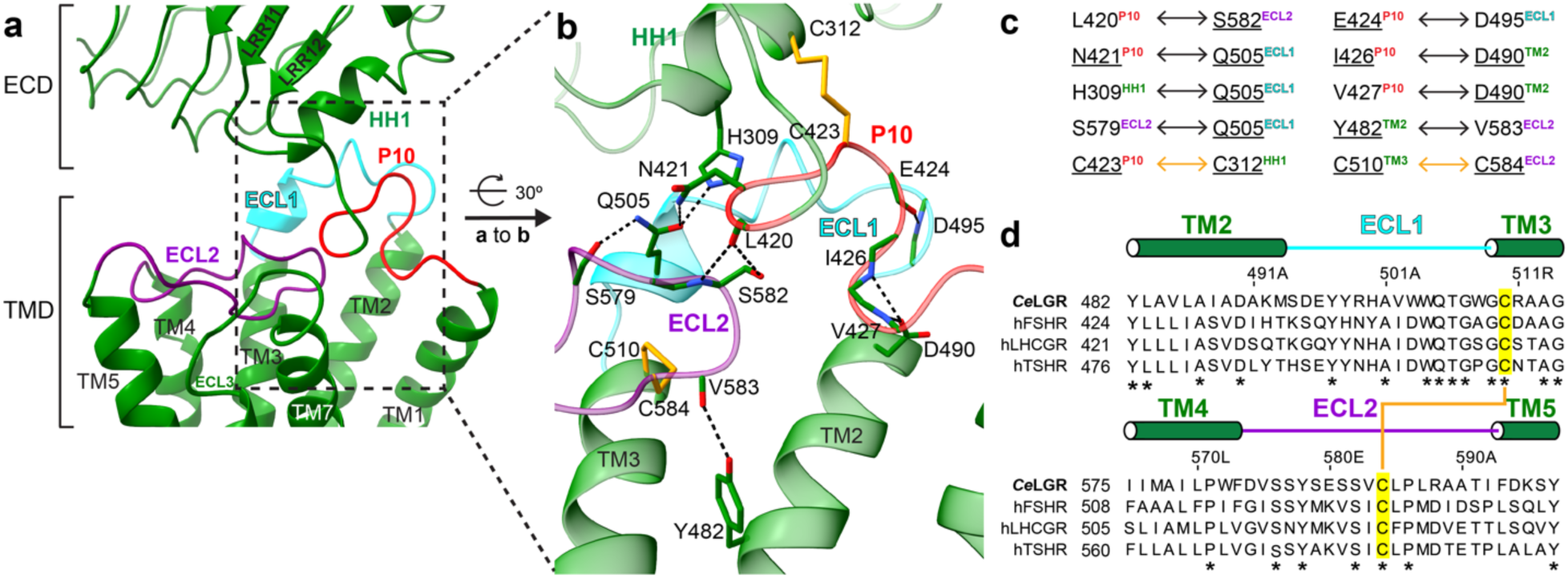
Interactions between the ECD and TMD of *Ce*LGR protomer A. **(a)** P10 in the CCR hinge region of *Ce*LGR plays a major role in linking the ECD with the TMD through interactions with HH1, ECL1, ECL2 and TM2. P10, ECL1 and ECL2 are colored in red, cyan and purple, respectively. **(b)** A magnified illustration of the region boxed in **a** to show atomic interactions between HH1, P10, TM2, ECL1 TM3 and ECL2. The ECL3 and TM7 are not shown for clarity. Hydrogen bonds are shown in black dashed lines. Disulfide bridges are represented as orange sticks. The carbon, oxygen and nitrogen atoms are shown in green, red and blue, respectively. **(c)** A summary of interactions shown in **b**. Conserved residues among *Ce*LGR and hGpHRs are underscored. Hydrogen bonds and disulfide bridges are indicated by black and orange arrow, respectively. **(d)** Sequence alignments in the vicinity of ECL1 and ECL2 comparing *Ce*LGR with the hGpHRs. Cysteine residues are highlighted with yellow background, and disulfide bridges are indicated by orange connections. The conserved residues are underscored by asterisks (*).

Residue Gln505 in ECL1 is essential in interaction with HH1, P10 and ECL2. The side chain of Gln505^ECL^^1^ forms hydrogen bonds with the side chain of His309^HH^^1^ and Asn421^P^^10^, and the carbonyl backbone of Ser579^ECL^^2^ (Fig. 3b). The ECL2 of *Ce*LGR interacts with TM2 through a hydrogen bond between the carbonyl backbone of V583^ECL^^2^ and side chain of Y482^TM^^2^, and TM3 through a disulfide bridge Cys584^ECL^^2^:Cys510^TM^^3^ (Fig. 3b and 3d).

The interactions mentioned above are summarized in Fig. 3c. Most residues involved are conserved between *Ce*LGR and all three hGpHRs (Fig. 2d and 3d). The ones which are not conserved (Leu420^P^^10^, Ser579^ECL^^2^, Val427^P^^10^ and V583^ECL^^2^) interact through backbone atoms; thus, variations at these residues will not affect their interactions. His309^HH^^1^ in *Ce*LGR has evolved to serine in all three hGpHRs (Fig. 2d). A systematic study of substitutions of Ser277 in hLHCGR, corresponding to His309^HH^^1^ in *Ce*LGR, indicated that the activity of S277H LHCGR did not change significantly from the wild-type receptor^41^.

In summary, the ECD and TMD of *Ce*LGR interact with each other through the CCR hinge region and extracellular loops in a delicate arrangement, which keep the pore to the TMD cavity ‘blocked’. The ECD of *Ce*LGR could be viewed as a tethered antagonist from this point.

### Dimer interface between *Ce*LGR protomers A and B

The *Ce*LGR protomers A and B form a compact homodimer through an extensive dimer interface that primarily involves their TMDs. In particular, TMs1, 6 & 7 from protomer A contact TMs1, 5, 6 & 7 from protomer B, with a buried surface area of 2075.2 Å^2^ from the TMDs (Fig. 4a to d). As mentioned earlier, the density map of TM6 and TM7 in both protomers is poor with no density observed for many side chains. The electron density for half of TM6 in protomer B could not be resolved at all, and it is indicated in grey in Figs. 4a to c. In addition, the central layers of direct homodimer contacts are flanked by an assigned cholesterol molecule, which facilitates the interaction between TM6 of protomer A and TM1 of protomer B (Fig. 4a to d).

**Fig. 4.**
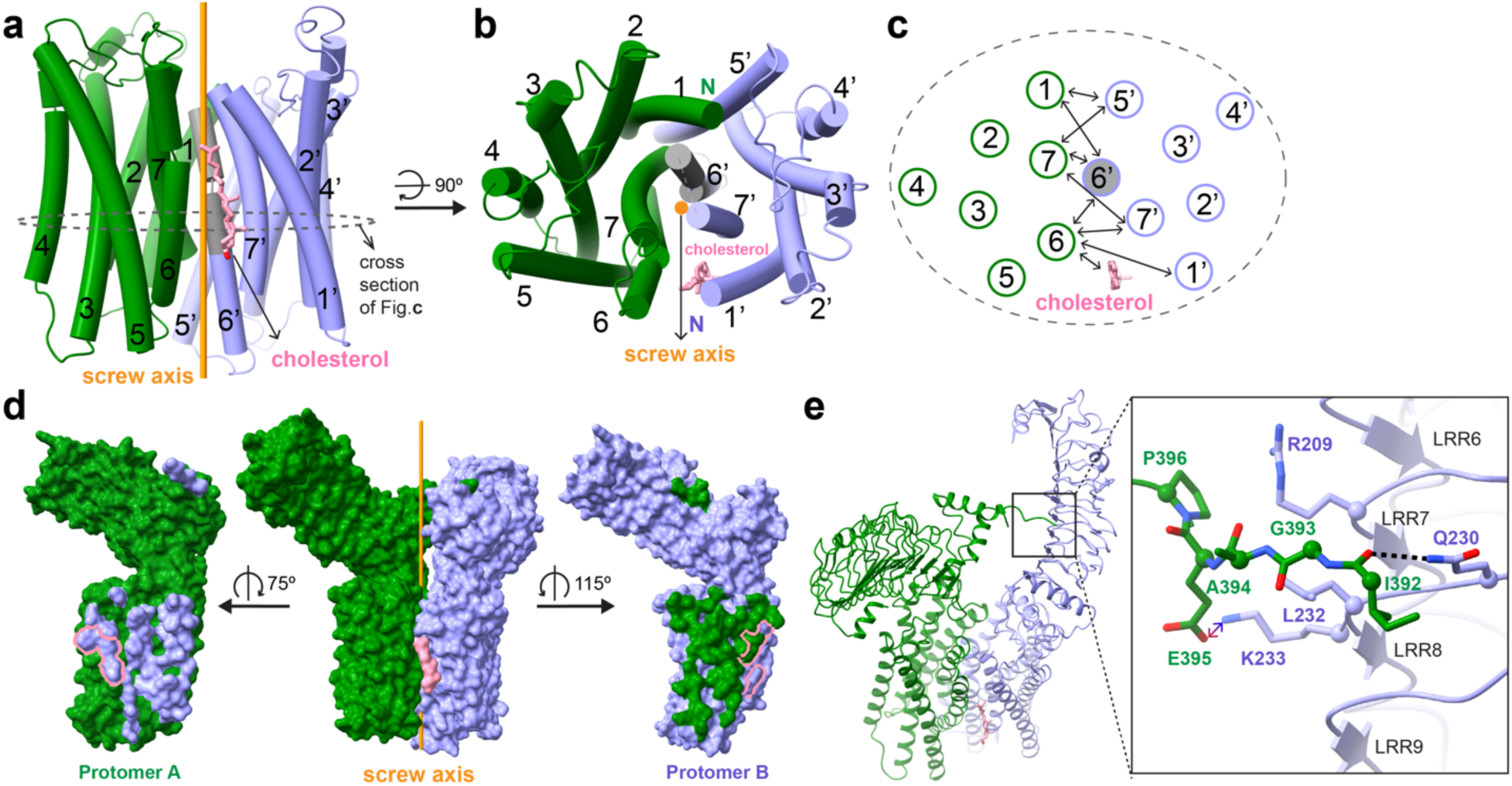
Dimer interface between *Ce*LGR protomers A and B. **(a)** Dimer interface between the TMDs of *Ce*LGR protomer A (green) and B (light purple). The transmembrane helices (TMs) are illustrated as cylinders and numbered. Half of TM6 in protomer B is shown in grey because its density could not be resolved for modeling. The orange stick is the screw axis from protomer B to protomer A. The cholesterol is shown as sticks (pink). **(b)** View of the TMD dimer interface from outside the cell. **(c)** View of a cross section of the homodimer as circled in **a**, but in the same view orientation as **b**. **(d)** Molecular surface of the homodimer (center) and then of protomer A (left) and protomer B (right) after opening by rotations as indicated and left with the imprint of the surface buried into the interface as colored by the partner protomers or by the assigned cholesterol outlined in pink. **(e)** Direct interaction between the CCR hinge region of protomer A and the ECD of protomer B. A hydrogen bond is shown as a black dashed line, and an ionic interaction is indicated by a double-headed arrow colored as for the contacting charged atoms.

Direct contacts are also observed between the CCR hinge region of protomer A and the ECD of protomer B (Fig. 4e) with a buried surface area of 185.3 Å^2^ . The backbone carbonyl group of Ile392 in protomer A forms a hydrogen bond with the side chain amino group of Gln230 in protomer B (Fig. 4e). The side chains of Glu395 in protomer A and Lys233 in protomer B make an ionic interaction (Fig. 4e). In addition, favored van der Waals contacts are observed between Leu232 in protomer B and Gly393, Ala394, Glu395 in protomer A, Gln230 in protomer B and Ile392 in protomer A, as well as Arg209 in protomer B and Pro396 in protomer A (Fig. 4e).

Urizar et al. have reported that hTSHR and hLHCGR form homo- and heterodimers via interactions involving primarily their TMDs and their ECDs are dispensable for dimerization. However, the ECDs must play a role in the dimeric interaction because differences were observed in the FRET signals obtained from dimers made of intact or truncated hTSHR devoid of ECD^8^. Our *Ce*LGR structure supports their observation that dimeric interactions involve primarily the TMDs. It is unlikely that the spare interactions between ECDs of the protomers (Fig. 4e) will affect receptor dimerization very much since the major dimer interface resides in the TMD; however, hormone binding may generate greater interaction.

### Conformational dynamics

Various features of the analysis indicate that *Ce*LGR in these preparations is highly dynamic. Distinct sub-populations could not be detected. Thus, we undertook a 3D variability analysis (3DVA) of the particles by *cryoSPARC*^42^, which corroborates and quantifies this impression (Supplementary Videos 1-2). The *Ce*LGR protomers appear to be in complex oscillatory motions about the interprotomer screw axis. Rotational orientations of the protomers vary smoothly through the trajectory, most obviously for the ECDs; each protomer contracts and extends along the screw axis; and the TMD micelle appears to expand and contract in diameter.

To further analyze these dynamics, we built models at the motional extrema by rigid body fittings of the ECD (from N terminus to LRR12) and TMD (TM1 to TM7) from each protomer into the initial and final frame of the computed 3DVA progression. These two models (conformers 1 and 2, respectively) were then compared with the model in the ‘rest’ state as built with the consensus frame (same with Fig. 1b). Protomers A and B fluctuate in opposite directions asymmetrically about the interprotomer screw axis (Fig. 5 a and b). As a result, relative to the rest state, the angle between the ECDs decreases by 6.9° for conformer 1 and increases by 11.1° for conformer 2. Similarly, the interprotomer angle for the TMDs decreases by 5.3° for conformer 1 and increases by 10.8° for conformer 2.

**Fig. 5.**
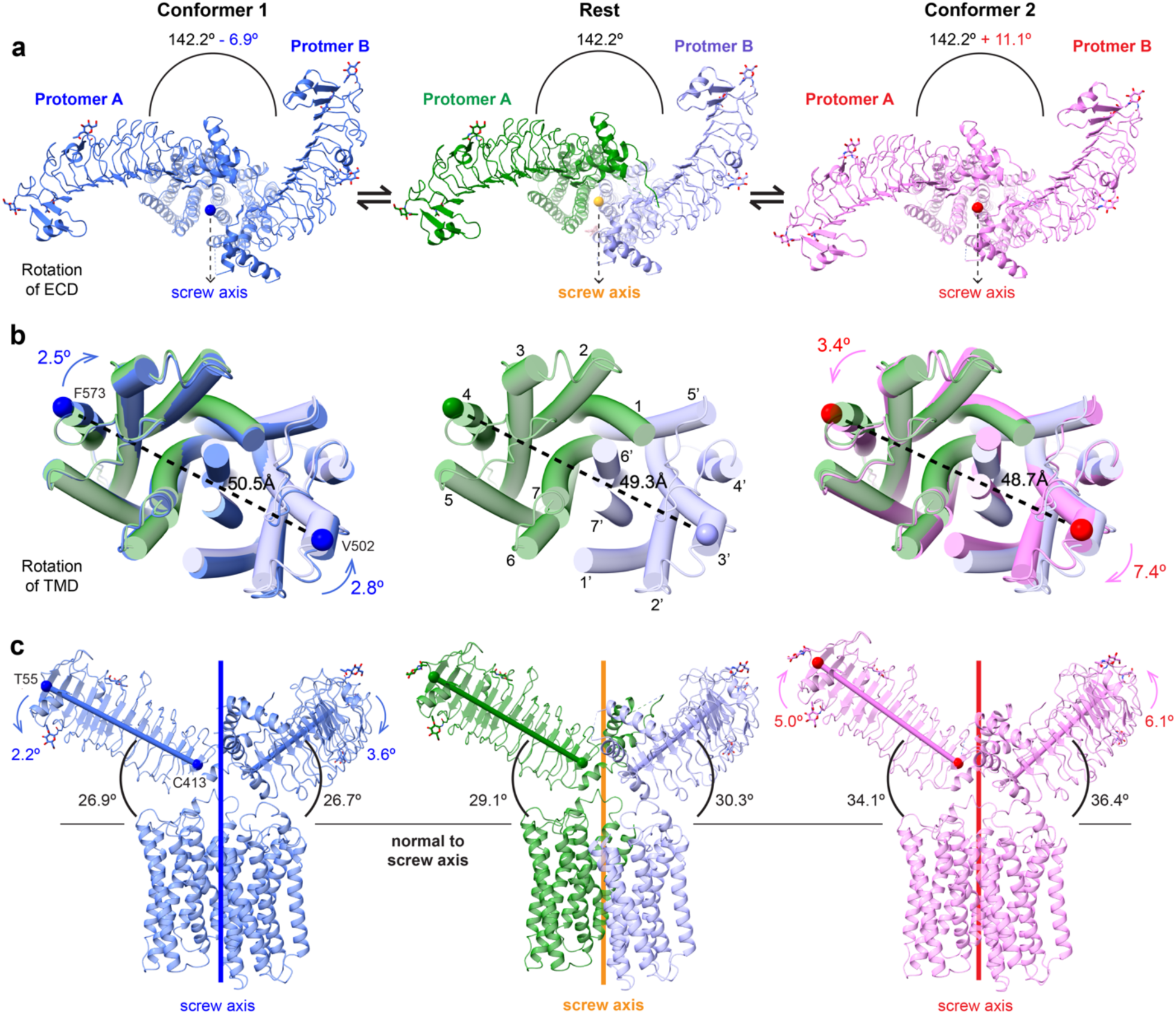
Fluctuational dynamics of the apo-state *Ce*LGR homodimer. The rest state model is in the center column (colored as in Fig. 1) and extreme conformers 1 (blue) and 2 (pink) are in the columns at left and right, respectively. (**a**) ECDs as viewed down the indicated screw axes with interprotomer rotations indicated. **(b)** TMDs as viewed into the membrane from extracellular side with TMs drawn as cylinders. The ‘rest’ state model (center) is superimposed on the extreme conformers (1, left; 2, right) with excursion rotations indicated by directed arrows. Measures of expansion and contraction are given as distances between markers Cα 573 in protomer A and Cα 502 in protomer B. **(c)** Side views of the *Ce*LGR homodimer as in Fig. 1b. The angles between the ECD axis indicator (Cα 55 and Cα 413) and the plane normal to the screw axis are given for each chain, and excursion rotations are indicated by directed arrows.

Concomitant with the rotations, the ECD of each protomer also moves relative to the deduced membrane surface (Fig. 5c). To measure this angle, we defined a pseudo axis from C_α_ 55 in β0 to C_α_ 413 in LRR12 and determined the angle between this axis and the plane normal to the screw axis. Taken again relative to the rest state, this angle decreases by 2.2° and 3.6° in protomers A and B, respectively, on going to conformer 1 and it increases by 5.0° and 6.1° in A and B, respectively on going to conformer 2. TMD positions relative to the membrane plane are relatively constant through the oscillations (Fig. 5c); however, screw dispositions of the TMDs do shift and they do so asymmetrically [(Δχ, Δt_χ_) = (−3.2°, - 0.5Å) and (+10.1°, - 1.6Å) respectively for conformers 1 and 2 relative to the rest screw (χ=140.0°, t_χ_=5.1Å) for TMD_B_ relative to TMD_A_]. At the same time, the TMD separations also vary (Fig. 5b); and, to quantify these changes, we define an outer surface width by the distance between C_α_ 573 in TM4 of protomer A and C_α_ 502 in TM3′ of protomer B. On going from conformer 1 to conformer 2, this distance decreases from 50.5 Å to 48.7 Å through the rest distance of 49.3 Å.

The extents of rotational displacements and ECD elevations are appreciably greater in protomer B than in protomer A. This is consistent with the observation that the density map quality is worse in protomer B than in protomer A. The extensive TMD interface is both intimate and highly dynamic, seeming to involve correlated movements between the two protomers.

### Activation of *Ce*LGR by *Ce*α2β5 *in vitro*

The *C. elegans* genome contains only one LGR sequence^29^, the type A LGR that is the subject of this study and only one LGR ligand, *Ce*α2β5^43^. It has recently been reported that the *Ce*α2β5:*Ce*LGR (called GPA2/GPB5:FSHR-1 in Ref. 10) pathway controls growth in *C. elegans*^11^. *Ce*α2β5 signaling is required for normal body size and intestinal lumens, acting through activation of *Ce*LGR for cAMP production^11^. In addition, *Ce*LGR (FSHR-1) has been implicated in responses to infection, oxidative and freezing-thawing stress in *C. elegans*^11^, which suggest that the thyrostimulin (alias for α2β5 hormone) signaling pathway may also regulate protective immune and stress responses in *C. elegans*.

To determine whether *Ce*α2β5 can activate *Ce*LGR, we applied a cAMP-based competitive assay in HEK293S cells. HEK293S cells infected with baculovirus carrying the *Ce*LGR gene were incubated with serially diluted *Ce*α2β5 hormone. The cAMP production was quantitatively determined to calculate the half maximal effective concentration (EC_50_) of *Ce*α2β5, which is 159.1 nM (Fig. 6a). The EC_50_ of recombinantly expressed and purified hTSH to wild type hTSHR was also measured as a control. The measured EC_50_ of recombinantly purified hTSH is 7.5 nM (Extended Data Fig. 8), which is comparable to that of the wild type hTSH (3 nM)^44^. Of interest, the G_s_ is highly conserved between human and nematode with sequence identity percentage of 66%^45^, consistent with the observed coupling between *Ce*LGR and G_s_ protein found in HEK293S cells. The potency of *Ce*α2β5 is lower than that of hTSH according to our measured EC_50_. The imperfect ‘match’ between *Ce*LGR and mammalian G protein, as well as the tight truncation of *Ce*LGR right after helix 8 may account for the lower potency of *Ce*α2β5.

**Fig. 6.**
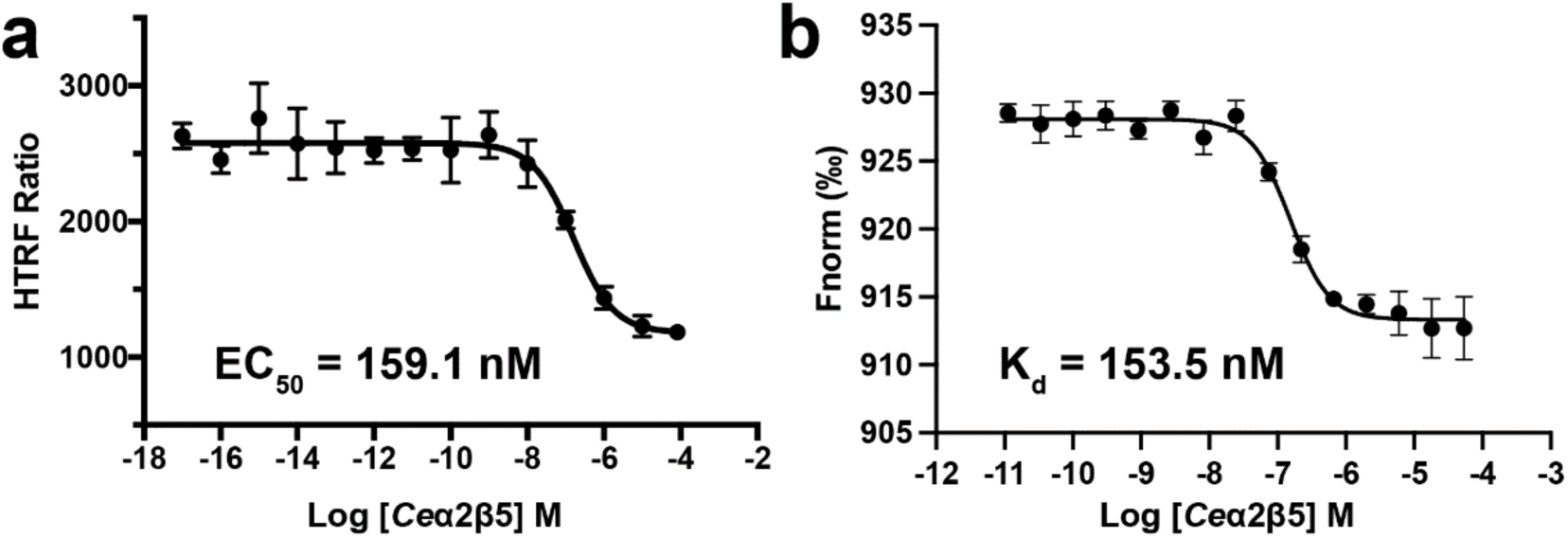
Activating interactions of *Ce*α2β5 with *Ce*LGR in vitro. **(a)** The EC50 for *Ce*α2β5 activation of *Ce*LGR was measured using a cAMP-based competitive assay in HEK 293S cells. The specific signal, homogenous time resolved fluorescence (HTRF) ratio, is inversely proportional to the concentration of native cAMP produced in the sample upon addition of *Ce*α2β5 ligand. Data points (mean ± s.d) are plotted for measurements from an experiment of n=3 biological replicates. **(b)** The equilibrium dissociation constant (Kd) of the *Ce*α2β5:*Ce*LGR complex was determined using MST. Data points (mean ± s.d.) are plotted for measurements from an experiment of n=4 biological replicates (see Methods for details).

In addition, the dissociation constant (K_d_) of the *Ce*α2β5:*Ce*LGR complex was determined as 153.3 nM using microscale thermophoresis (MST) (Fig. 6b), which is in a comparable range to the EC_50_ of *Ce*α2β5.

## Discussion

This study reveals several structural features of the LGR from *C. elegans* that are of interest in relation to the functioning of this particular receptor and, by analogy, to the activity of related receptors. Most prominently, this nearly full-length LGR is dimeric, both in solution after detergent extraction and as imaged on cryo-EM grids. These dimers show pronounced asymmetry, and they are highly dynamic in the apo state, without the cognate α2/β5 hormone and heterotrimeric G protein being present. Moreover, the receptor chains have been cleaved in the CCR hinge region at a site between disulfide-bridged cysteine residues.

### *Ce*LGR in relation to mammalian glycoprotein hormone receptors (GpHRs)

*Ce*LGR and the mammalian GpHRs are similar in sequence and in structure, and the relationship is closest to TSHR. The co-evolution of type A LGRs and their activating ligands is thought to have proceeded in early vertebrates in two rounds of whole genome duplication from invertebrate counterparts, the first producing progenitors of TSH and TSHR and the second adding FSH and LH and their receptors; thus, by evolution, *Ce*LGR is expected to be most closely related to TSHR. Indeed, *Ce*LGR is like hTSHR in having a sizable insertion, relative to hFSHR and hLHCGR, within its CCR hinge region and, also, in being cleaved just beyond HH2 between the disulfide-bridged cysteine residues of the CCR (Extended Data Fig. 1). As for *Ce*LGR, helices HH1, HH2 and HH3 are all ordered in structures of TSHR^4,5^, whereas only HH1 is ordered in the structures of LHCGR^3^ and FSHR^6^. Additionally, just as *Ce*LGR binds *Ce*α2β5, human TSHR uniquely retained high affinity for *Hs*α2β5 (thyrostimulin) while it acquired affinity for hTSH (thyrotropin) as well. Finally, the asymmetric *Ce*LGR dimer structure is consistent with abundant evidence for functional GpHR dimerization^7,8,10,16–24^, which is particularly compelling for hTSHR^8,10,16,20,21^.

The *Ce*LGR dimer differs from other GpHR dimers in the strength of its self-association. Whereas we found no monomers in our 2D classifications of cryo-EM particles, and only a small fraction (1.5%) of trimers (Extended Data Fig. 4), only monomer classes were reported for the cryo-EM analyses of human GpHRs^3–6^. Monomers also predominate (70%) in imaging analyses of FSHR as expressed in HEK293 cells; however, a substantial fraction of dimers (15.5%) and other oligomers are also present in basal conditions, with some variation in the presence of hormone variations^46^. Similar oligomerization levels (41.4%) were found for LHR expressed in HEK293 cells^19^. Of course, dimerization is concentration dependent; and, in the case of TSHR, the expression level in thyrocyte cultures needed to be at the thyroid level *in situ* to exhibit physiological biphasic responses reflective of dimerization^9^.

The puzzle as to why detergent-extracted *Ce*LGR is dimeric whereas similarly purified human GpHRs are monomeric may be explained by properties of the dimer interface of *Ce*LGR. Nearly all of the surface area buried into that interface comes from the transmembrane helices (91.8%), for which the sequences are highly conserved among the three human GpHRs (68-72% identity overall) and especially so for residues at the TMD interior (84-88% identity). The respective identity levels between the *Ce*LGR and human GpHR sequences, while appreciably lower, are also quite high (35-37% overall and 58-63% for interior residues). The dimer-interface residues of *Ce*LGR are more conserved than outward-facing residues generally (29-37% identity with human GpHR homologs as compared to 25-28% overall). Similarly, among human GpHRs, the homologs of dimer-interface residues are more conserved than are all outward-facing residues (63-74% compared to 61-67% identity overall). The *Ce*LGR interface includes seven residues that are distinguished from counterparts that are identical in human GpHRs and in *Drosophila* LGR1 as well (A442L, T637K, V644F, L647F, V675L, F685C and V692A). As for the human GpHRs, detergent-extracted *Dm*LGR1 purifies as a monomer (data not shown). We postulate that the *Ce*LGR interface distinctions stabilize these dimers, whereas the similar but distinctive interfaces of the homologous receptors are more labile and do not survive detergent solubilization from the lipid bilayer.

### The *Ce*LGR dimer as related to dimerization of other GPCRs

Whether the *Ce*LGR structure truly reflects the nature of functional dimers of TSHR and other GpHRs remains to be seen. Meanwhile, since the evidence for physiological relevance of GpHR dimers is compelling, the *Ce*LGR dimer provides us with hypotheses on the functioning of GpHRs. The growing body of evidence on the obligate dimers of class C and class D GPCRs^47^ supplements this structural foundation. In particular, functional complementation such as that employed in GpHR experiments^7,8,21–24^ is intrinsic to GABA_B_ receptors, which are heterodimeric class C GPCRs wherein the B1 subunit binds the ligand but does not contact Gα and the B2 subunit does not interact with GABA but does bind Gα^25,26^. The homodimeric class C GPCRs, metabotropic glutamate receptor^48^ and calcium-sensing receptor (CaSR)^49^, also have only one G protein coupled to the receptor dimer.

The TMD interfaces in the known class C structures are more symmetric than that in the *Ce*LGR structure; but they nevertheless involve similar TMD surfaces. For the active GABA_B_ and CaSR receptors, these are centered at TM6-TM6′ interactions with only peripheral involvement of the adjacent TM5 and TM7 helices; for *Ce*LGR, being off center, TM6 and TM7 of protomer A contact the TM5-TM7 surface. Also, whereas direct contacts are appreciable in the active states for class C receptors, and they vary but are typically minimal in inactive states^25,48,50,51^, the *Ce*LGR dimer interface in the hormone-free (presumably inactive) state here is intimate albeit mobile. Such TM6-centered dimers may not be universal, however; cross-linking evidence for 5HT2c and some other class A receptors contacts involving TM4 and TM5^52^.

### Implications for receptor activation

The structure of the *Ce*LGR dimer sheds light on the activation mechanism of GpHRs. Its asymmetric assembly suggests a structural basis for the negative cooperativity observed in hormone binding^8,20^, for its corollary in switching from Gs signaling from 1:2 complexes to Gq or Gi/o signaling in 2:2 complexes^9,10,21^, and for the functional complementation provided by the co-expression of binding-deficient and signaling-deficient receptors^7,8,22–24^. The current activation model based on structures of monomeric GpHRs^3–6^ could not explain these effects, whereas the compactly associated *Ce*LGR dimer makes allosteric modulation and rescue of signaling possible. Atomic details of the conformational changes found in the full-length GpHR monomers upon binding of their hormones coupled with constraints from the dimer structure allow us to contemplate how this might happen.

As found for the hTSHR, hFSHR and hLHCGR monomers in their inactive hormone-free states, both *Ce*LGR protomers have their ECDs in a ‘down’ orientation. Upon activation for hormone binding, each GpHR then adopts an ‘up’ conformation with its ECD rotated upward by 45-55° relative to the ligand-free state^3–6^. We presume that *Ce*LGR similarly adopts an ECD-up conformation when hormone associated and that the *Ce*LGR asymmetry is retained as it forms a 1:2 complex. Whether such dimers have both protomers ECD-Up or only the one that is hormone-bound is uncertain; however, we successfully modeled both alternatives. We built *Ce*α2β5-activated *Ce*LGR dimers by first superimposing the TMD of hTSHR in a hTSH:hTSHR:G_s_ complex (PDBid 7UTZ) onto each *Ce*LGR protomer, then moving one or both of the *Ce*LGR ECDs into superposition with the ECDs of this TSHR ‘dimer’, and finally replacing hTSH on protomer A with the *Ce*α2β5 ligand^27^ (Extended Data Fig. 9a).

*Ce*α2β5 could only be fitted to protomer A because the binding site in protomer B, whether ECD-up or ECD-down, is blocked by the CCR hinge region of protomer A as it is for both ECDs down (Fig. 4e). As noted for hGpHRs^3–6^, *Ce*α2β5 would be expected to collide with the cell membrane if bound to *Ce*LGR in the down conformation (Extended Data Fig. 9b). Moreover, there is no room for a second *Ce*α2β5 unless the subunits are rotated, with both ECDs up, from asymmetry (142° apart) to symmetry (180°). In accord with the observed negative cooperativity, this symmetrization entails an energetic penalty for the second-site binding event, which is lessened when the receptor is pre-disposed by activating mutation^20^ to having both protomer ECD-up for the 2:2 state.

Another question is how many heterotrimeric G proteins couple to a *Ce*LGR dimer? There is sufficient room for only a single heterotrimeric G protein at a time to bind to the 1:2 *Ce*LGR dimer; however, G-protein coupling with either protomer A (cis activation) or B (trans activation) seems possible (Extended Data Fig. 9a). In light of functional complementation experiments for hGpHRs and structural results from class C GPCRs, trans-activation would be expected. The actual activation mechanism for dimeric *Ce*LGR is complicated and remains to be determined experimentally.

### Implications for physiology

Little is known about the molecular physiology of the LGR-α2β5 system in nematodes, but much is known about the GpHR-hormone systems of humans. What role the dimer may play in the observed impact of the LGR system in worm physiology, and whether the tight association in the *Ce*LGR dimer is important for its functioning is unknown. As discussed above, there is considerable evidence for the role of GpHR dimers in human physiology; however, these dimers are relatively loosely self-associated since they do not survive detergent-extraction from cell membranes. Moreover, TSHR dimerization in cells requires expression at the level of thyrocytes *in vivo*^9^. How labile GpHR dimerization impacts biological function is unclear; but the resulting effects of negative cooperativity are evident: whereas the 1:1 hormone:dimer complex promotes Gs signaling and cAMP production, the 2:1 complex from higher TSH levels promotes Gi/Go signaling to elicit IP1 production and a biphasic cAMP response^9,10,21^.

Negative cooperativity, resulting from allosteric communication between protomers of the dimer, is essential for the function of GpHRs. For instance, while the hCG level is very low (0.04 to 5.5 ng/ml) for unpregnant women, it rises rapidly during pregnancy and can reach 1,952 to 19,958 ng/ml at a peak around 8-10 weeks^53^. Negative cooperativity provides exquisite sensitivity at low hormone concentrations while buffering against acutely elevated hormone levels^54^.

## Materials and methods

### Homolog screen and construct design

Homologs of GpHR from the following species were cloned into a BacMam vector with oxidizing environment-optimized GFP (oxGFP)^55^ fused at the C-terminus for fluorescence-detection size exclusion chromatography (FSEC) screen^56^: *Caenorhabditis elegans* (UniProt code: G5EG04)*, Drosophila melanogaster* (Fruit fly; UniProt code: Q9VEG4)*, Petromyzon marinus* (Sea lamprey, UniProt code: Q0P6K8)*, Danio rerio* (Zebrafish; UniProt code: F1R4X9)*, Xenopus laevis* (African clawed frog; UniProt code: G9M6I0)*, Bothrops jararaca* (Jararaca; UniProt code: Q6K0L4)*, Gallus gallus* (Chicken; UniProt code: P79763)*, Rattus norvegicus* (Rat; UniProt code: P20395)*, Mus musculus* (Mouse; UniProt code: P35378)*, Bos taurus* (Bovine; UniProt code: P35376) *and Homo sapiens* (Human; UniProt code for hFSHR: P23945, hLHCGR: P22888 and hTSHR: P16473). The codons from each species other than human have been optimized for expression in human cells. An influenza hemagglutinin (HA) signal peptide was fused at the N terminus of each gene in place of its original signal peptide to enhance protein insertion into cellular membranes. *Ce*LGR (UniProt accession code: G5EG04) was identified as the best candidate for structural studies because of its large expression yield and good solubility in detergent.

*Ce*LGR contains a large intracellular domain (Ile713 to Ser929) at its C-terminus compared with hGPHRs (Extended Data Fig. 1), which is predicted by AlphaFold to be unstructured. In order to reduce structural flexibility potentially introduced by the unstructured C-terminus, *Ce*LGR was truncated after helix 8. Therefore, the final construct contains an HA signal peptide replacing the natural signal peptide (*Ce*LGR residues 1 -31) at the N-terminus, followed with a Flag tag, residues Gln31 to Arg712 of *Ce*LGR, a triple alanine linker and a Flag tag at the C-terminus for affinity purification. The oxGFP was removed by inserting a stop codon after the C-terminal Flag tag. The presence of both N-terminal and C-terminal Flag tag is because a proteolytic posttranslational cleavage was observed during the maturation event of *Ce*LGR. The original plan was to purify the N-terminal Flag-tagged ECD with anti-Flag M1 antibody affinity resin and the C-terminal Flag-tagged TMD with anti-Flag M2 antibody affinity resin. Actually, the cleaved ECD and TMD of *Ce*LGR are linked by three disulfide bridges in the CCR hinge region. Therefore, either one of the two Flag tags should be sufficient to purify the *Ce*LGR.

### Expression and purification of *Ce*LGR

The gene of the expression construct was cloned into a BacMam expression vector containing a human cytomegalovirus (CMV) promoter. Baculovirus was made in Sf9 cells. For large-scale expression, suspension-adapted HEK293S, which lack N-acetyl-glucosaminyltransferase I (GnTI), were grown in Freestyle 293 expression medium (Life Technology) supplemented with 2% Fetal Bovine Serum (FBS) at 37 °C in the presence of 8% CO_2_. The culture was transduced with P3 baculovirus once cell density reached 2.0 x 10^6^ to 3.5 x 10^6^ cells per milliliter. After incubation for 8-16 h at 37 °C, 10 mM sodium butyrate was added to enhance the expression level, and the temperature was changed to 30 °C for protein expression. Cells were harvested 72 h after adding sodium butyrate and resuspended in a buffer containing 20 mM HEPES, pH 7.5, 300 mM NaCl and 10 mM MgCl_2_, supplemented with EDTA-free protease inhibitor cocktail tablet (Sigma) and 1 mM phenylmethylsulfonyl fluoride (PMSF). Cell pellets were stored at -80 °C until use or lysed immediately for protein purification.

Cells were lysed using an EmulsiFlex-C3 high pressure homogenizer (Avestin) with three passes at 10,000-15,000 psi. Cell debris was removed by centrifugation at 2850 x g for 15 min. Cell membranes were collected by ultracentrifugation for 1 h in a Beckman 45 Ti rotor at 40,000 r.p.m. The membranes were homogenized and subsequently solubilized for 2 h in a buffer composed of 20 mM HEPES, pH 7.5, 150 mM NaCl, 10 mM MgCl_2_, 1% lauryl maltose neopentyl glycol (LMNG), and 0.2% cholesteryl hemisuccinate tris (CHS), supplemented with protease inhibitor cocktail tablet (Sigma) and 1 mM (PMSF). After insoluble material was removed by ultracentrifugation for 1 h at 32,000 r.p.m. in a Beckman 50.2 Ti rotor, the supernatant was incubated with anti-FLAG M2 resin (Sigma) by rotating overnight. The resin was washed with 30 column volumes of wash buffer containing 50 mM Tris HCl, pH 7.4, 150 mM NaCl, 0.01% LMNG and 0.002% CHS and the protein was eluted using wash buffer supplemented with 0.2 mg/ml Flag peptide (Sigma). The eluted protein was concentrated using a 100-kDa cutoff concentrator and injected into a Superose 6 Increase size exclusion column (Cytiva) equilibrated in a buffer composed of 20 mM HEPES, pH 7.5, 150 mM NaCl, 10 mM MgCl_2_, 0.002% LMNG, and 0.0004% CHS. The peak fraction was collected for imaging. All purification steps were conducted on ice or at 4 °C.

### Construct design, expression and purification of hTSHR

The wild type hTSHR (UniProt code: P16473, residues 21-764) was cloned into a BacMam expression vector containing a CMV promoter with an HA signal peptide at its N-terminus in place of its native signal peptide and a Flag tag at its C-terminus. The expression and purification procedures for hTSHR were the same as those used for *Ce*LGR.

### Characterization of cellular proteolysis of expressed *Ce*LGR and hTSHR

The purified *Ce*LGR and hTSHR receptors were analyzed by SDS polyacrylamide gel electrophoresis (SDS PAGE) with or without the presence of a reducing agent DTT to break the disulfide bridges. To test whether residual monomer bands reflected intact receptors or other effects, the purified proteins also went through limited trypsin digestion by adding trypsin to a final concentration of 0.005% to 0.01% followed with incubation at room temperature for 15 min. A 2-fold excess of trypsin inhibitor (Sigma) was then added, and incubation continued for 10 min before adding DTT.

The site of natural protease cleavage was delimited by mass spectrometric (MS) analysis of proteolytic peptides after digestion of the denatured receptors. Because natural cleavage was incomplete for the receptors as expressed in HEK293S cells, we used proteolytic MS analysis of only the post-cleavage bands from SDS PAGE (highlighted in red boxes in Extended Data Fig. 2). The gel band was excised, and *in-gel* digestion was performed as previously described^57^ with minor modifications. Briefly, the proteins were digested using trypsin, chymotrypsin or GluC. The resulting peptides were dissolved in a solution of 3% acetonitrile and 0.1% formic acid. Peptides were separated using a reversed-phase C18 column (25 cm × 75 µm, 1.6 µm, IonOpticks). Peptide MS/MS analysis was performed using a Thermo Scientific™ Orbitrap Fusion™ Tribrid™ mass spectrometer.

Raw mass spectrometric data were analyzed using the MaxQuant environment v.2.6.1.0, utilizing Andromeda for database searching with default settings with a few modifications. The default tolerances were used for the first search (20 ppm) and the main search (6 ppm). MaxQuant was configured to search against the reference human/*C. elegans* proteome database, which was downloaded from UniProt. The search parameters included trypsin/ chymotrypsin /GluC digestion with up to 2 missed cleavages. The false discovery rates (FDRs) for peptides, sites, and proteins were all set to 1%. The output combined folder was uploaded into Scaffold 5.0 (Proteome Software). Spectral counting was employed for comparative sample analysis.

### Expression and purification of *Ce*α2β5 and hTSH

*Ce*α2β5 was expressed and purified as previously described^12^ with a brief summary here. The genes coding for *Ce*α2 (UniProt code: A0T3A2, residues 29-120) and *Ce*β5 (UniProt code: A7DT38, residues 20-125) were cloned into two BacMam expression vectors. Both *Ce*α2 and *Ce*β5 have an N-terminal HA signal peptide in place of the original signal peptide. A Flag affinity tag was engineered between the HA signal peptide and *Ce*α2, while an HA affinity tag was engineered between the HA signal peptide and *Ce*β5 to facilitate affinity purification. Suspension-adapted HEK293S (GnTI^-^) cells were used for co-expression of *Ce*α2β5 using baculovirus produced by Sf9 cells. The *Ce*α2β5 protein was purified using anti-Flag M2 antibody affinity column and size exclusion chromatography.

The hTSH construct comprises the native signal peptide from hTSHβ (UniProt code: P01222, residues 1-20), followed by an HA affinity tag, the mature sequence of hTSHβ (UniProt code: P01222, residues 21-138), a 15-residue linker (GGGSGGGSGGGSGGG), the mature sequence of hα (UniProt code: P01215, residues 25-116), a triple alanine linker and a Flag affinity tag at C-terminus, which is analogous to the construct used by Fan and Hendrickson for the structural determination of hFSH bound to the extracellular hormone-binding domain of its receptor using X-ray crystallography^13^. Suspension-adapted HEK293S (GnTI^-^) cells infected with baculovirus secreted the hTSH into the cell medium. Both anti-HA (Sigma) and anti-Flag M2 antibody affinity resins were tested to purify the hTSH. No protein could be purified using the anti-HA affinity resin. It is very likely that the anti-HA resin could not bind to HA-tagged proteins well (not limited to hTSH) according to our experience. Finally, the hTSH was purified using an anti-Flag M2 antibody affinity column and Superdex 75 Increase size exclusion chromatography.

### Cryo-EM sample preparation and data collection

EM grids (Quantifoil R0.6/1, 400 mesh, holey gold) were glow discharged for 25 s (6.4 sccm H_2_, 27.5 sccm O_2_, 10 watts) using a Gatan Solarus model 950 Advanced Plasma Cleaning System prior to sample application. 3 ul of protein solution at 0.3-0.4 mg/ml was applied onto a grid and then plunge-frozen in liquid ethane cooled by liquid nitrogen using a Vitrobot Mark IV (FEI). The Vitrobot chamber was set to 100% humidity at 4 °C. The sample was blotted for 8 s with a wait time of 30 s and a blot force of 3.

Cryo-EM imaging was performed on a Titan Krios electron microscope (FEI) operating at 300 kV, equipped with a post-column GIF Quantum energy filter with a slit width of 20eV, and a K2 summit direct electron camera (Gatan) in counting mode. Micrographs were collected with 70 μm C2 aperture at a calibrated magnification of 130,000 corresponding to a magnified pixel size of 1.06 Å. Each micrograph comprises 50 sub-frames with a total accumulated dose of 71.01 e^-^/Å2, recorded for a total exposure time of 10 sec at a dose rate of 8 e^-^/pixel/sec. Data acquisition was done using Leginon across a nominal defocus range of -1.2 μm to -2.0 μm. A total of 8,028 micrographs were acquired as does-fractionated image stacks.

### Cryo-EM image processing

The cryo-EM data processing workflow is summarized in Extended Data Fig. 4. Maps and raw videos have been deposited at EMDB (Extended data Table 1) and EMPIAR (EMPIAR-11917). Subsequent steps were performed in cryoSPARC (v4.5.0) unless otherwise indicated^58^.

Patch-based motion correction and dose weighting of the 8,028 cryo-EM movies were executed in cryoSPARC using the Patch Motion job type. Patch-based CTF estimation was conducted on the aligned averages utilizing Patch CTF. Out of these, 7,031 micrographs were selected based on a CTF estimate resolution of < 4 Å, relative ice thickness between 0.9-1.1, total motion distance of < 50 pixels, and a defocus range of < 1200.

The initial round of particle picking employed Blob Picker, extracting particles in a 256-pixel box. Six typical classes emerged post 2D classification. The secondary round of particle picking used template picking with the six identified classes, resulting in 5,634,393 particles being picked. Subsequent multiple rounds of 2D classification yielded 997,598 selected particles. One well-defined class, consisting of 185,961 particles, was identified and selected after 3D classification. Nonuniform refinement of this class achieved a resolution of 3.87 Å. For the third-round particle picking, 75 exemplary micrographs, containing 70-90 good particles each (totaling 5,538 particles), were selected as the training dataset.

A tertiary round of particle picking utilized Topaz^59^, initially training the Topaz neural network with the 75 micrographs in the training dataset. In total, 965,789 particles were selected using the Topaz training model. Post 2D classification, 940,614 particles were chosen, with one well-defined class being selected after 3D classification. Subsequently, 286,046 particles corresponding to *Ce*LGR were re-extracted with re-centering in a 256-pixel box.

An initial refinement of all 286,046 *Ce*LGR particles, using nonuniform refinement, yielded a consensus refinement with a resolution of 3.82 Å. The application of dynamic masking and real-space windowing of particle images during refinement was not enabled. Further 3D classification without alignment resulted in the best class with complete ECD and TMD, consisting of 72,220 particles, achieving a nonuniform refinement resolution of 3.79 Å.

### 3D variability analysis

After 3D classification, 3D Variability Analysis^42^ (3DVA) was carried out with 286,046 selected particles in cryoSPARC, solving in three modes. Reconstructions were calculated along each mode. A mode corresponding to rotation of each protomer, in combination of contraction and extension of the *Ce*LGR dimer was identified. This mode was then used (using the 3D Variability Display job type) to split the particles into 20 clusters, and a reconstruction calculated for each to display the 3D variability.

### Atomic model building and refinement

An initial model for the *Ce*LGR homodimer, which includes residues from the N-terminus to TM5 in each protomer, was generated using phenix.predict_and_build^60^. This model was then extended manually and completed in Coot^61^. The cholesterol molecule and glycans were added where justified by the density map and chemical environment. The model was finalized by rebuilding in ISOLDE^62^ and refinement with phenix.real_space_refine^63^. Figures were prepared using ChimeraX^64^. Refinement and validation statistics are provided in Extended Data Table 1.

### cAMP assay

The same construct for expression of *Ce*LGR and hTSHR was cloned into a pcDNA3.1 vector for the cAMP assay. HEK293S (GnTI^-^) cells grown in Freestyle 293 expression medium supplemented with 2% FBS were seeded onto a 6-well plate at 1.0 x 10^6^ cells per milliliter. Two hours later, cells were transiently transfected with hTSHR plasmids using Lipofectamine 3000 (Invitrogen) or infected with P2 virus of *Ce*LGR, the same construct used for protein preparation for structural analysis. Transiently transfected (hTSHR) or virus-infected (*Ce*LGR) cells were incubated at 37 °C in the presence of 8% CO_2_ for two days. Next, cells were harvested with centrifugation at 200 xg for 5 min, washed with phosphate-buffered saline (PBS), resuspended in stimulating buffer (Cisbio), seeded onto a 384-well plate and incubated with serially diluted ligand for 2 h at 37 °C in the presence of 8% CO_2_. The cAMP accumulation was measured using the cAMP G_s_ dynamic kit (Cisbio) following manufacturer’s instructions. In this competitive assay, cAMP is covalently labeled with a red-emissive acceptor dye d2, and an anti-cAMP antibody is labeled with a cryptate donor. A Förster Resonance Energy Transfer (FRET) signal is detected when the d2-labeled cAMP (acceptor) binds to an cryptate-labeled anti-cAMP antibody (donor)^65^. Upon stimulation of *Ce*LGR in HEK293S cells with addition of *Ce*α2β5, native cAMP is generated. After cell lysis and the addition of two labeled compounds, unlabeled native cAMP will compete with the d2-labeled cAMP, and thus a decrease of FRET is observed. The specific signal is inversely proportional to the concentration of native cAMP produced in the sample. The fluorescence emissions at two different wavelengths (665 nm and 620 nm) were read on the EnVision 2104 Multilabel Microplate Reader (Perkin Elmer). Data were analyzed using GraphPad Prism 7.0 and were presented as mean ± standard deviation (s.d.) of measurements from an experiment of n=3 biological replicates.

### MST measurements

The dissociation constant (K_d_) of *Ce*α2β5 to *Ce*LGR was measured using Microscale Thermophoresis (MST). *Ce*α2β5 and *Ce*LGR were purified separately as described above. Purified *Ce*LGR was labeled with the RED fluorescent dye NT-647-MALEIMIDE using the Monolith NT Protein Labeling Kit RED - MALEIMIDE (NanoTemper Technologies) following manufacturer’s instructions. 50 nM of labelled *Ce*LGR dimer was incubated with serially diluted *Ce*α2β5 with a 1:1 (v:v) ratio. The normalized fluorescence (F_norm_) of the mixture was read on the Monolith NT.115 instrument (NanoTemper Technologies) selecting LED color in red. Data were analyzed using GraphPad Prism 7.0 and were presented as mean ± s.d. of measurements from an experiment of n=4 biological replicates.

### Data availability

The cryo-EM density map for *Ce*LGR in the apo state have been deposited in the Electron Microscopy Data Bank under accession code EMD-43735. Atomic coordinates for the *Ce*LGR structure in apo state have been deposited in the RCSB Protein Data Bank under accession code 8W1Z. Raw cryo-EM images have been deposited in the Electron Microscopy Public Image Achieve (https://www.ebi.ac.un/pdbe/emdb/empiar) under accession code EMPIAR-11917.

## Acknowledgements

Cryo-EM data were collected at Columbia Cryo-EM facility and the Simons Electron Microscopy Center (SEMC), with the assistance of staff from both SEMC and the Columbia University Cryo-Electron Microscopy Center. R. Grassucci and Z. Zhang from the Columbia Cryo-EM Center assisted with data collection. Some of the work was performed using equipment from the Center for Membrane Protein Production and Analysis (COMPPÅ; grant no. NIH P41 GM116799 to W.A. Hendrickson). We thank Dr. R. Bruni for the gift of BacMam-oxGFP vector. Z. Gong is partially supported by Endocrinology Training grant from NIH (grant number 5 T32 DK 7328-40).

## Author contributions

Z.G., Q.R.F. and W.A.H. designed the expression constructs. Z.G. performed gene cloning, protein purification, cAMP assay and MST measurement. B.K. and Z.G. carried out FSEC screens. S.C., Z.F. and Z.G. screened and optimized sample vitrification, and generated cryo-EM data. Single-particle cryo-EM data analysis was conducted by S.C., O.B.C., C.W., and Z.G. 3D Variability Analysis was carried out by Z. G. Model building was conducted by Z.G. The project was supervised by Q.R.F. and W.A.H. The manuscript was written by Z.G. and W.A.H with input from S.C., O.B.C., and Q.R.F.

---

**Extended Data Fig. 1.**
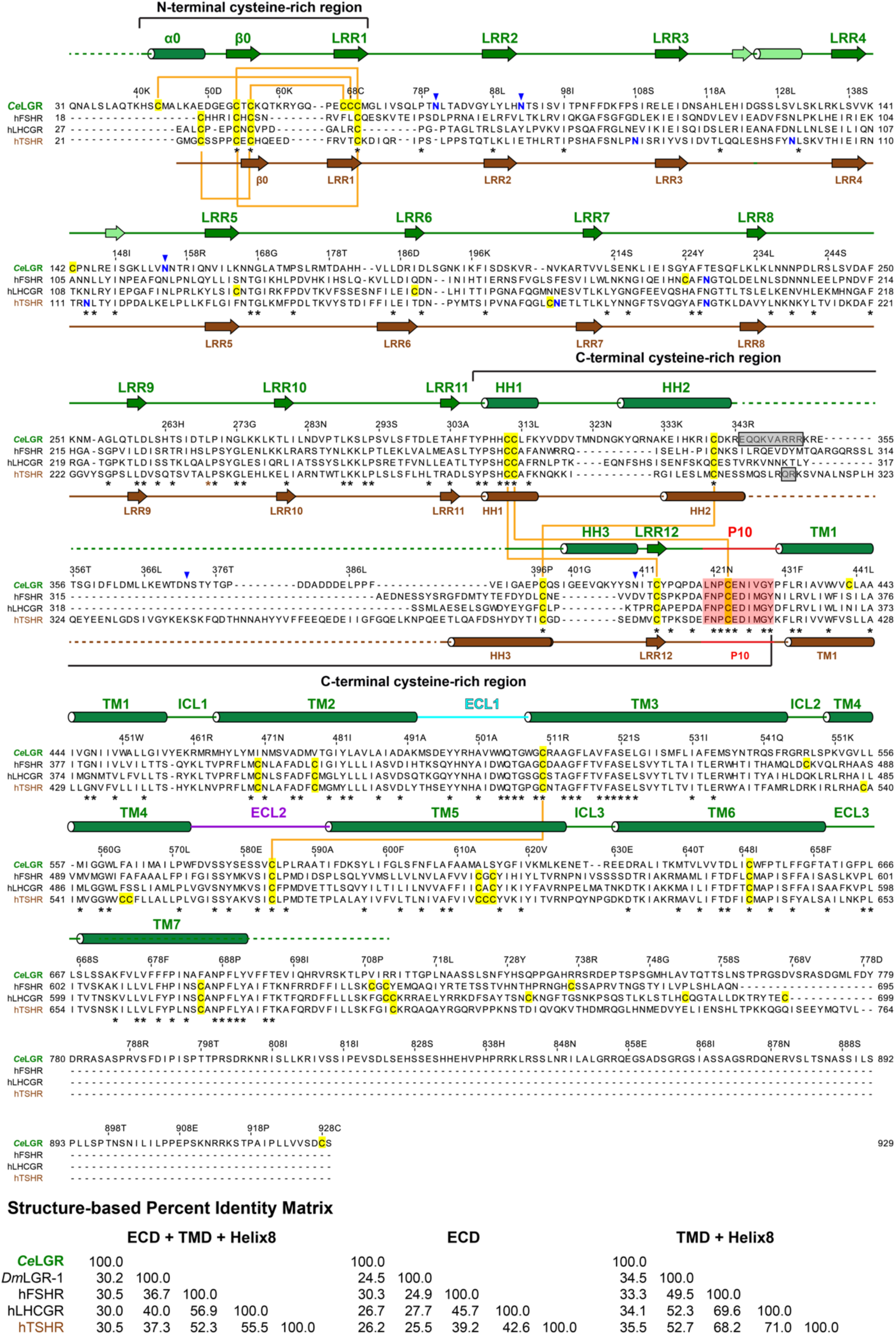
Structure-based sequence alignment of LGRs. The secondary structural elements are indicated by cylinders for α helices and arrows for β strands, green for *Ce*LGR and brown for hTSHR. Strands at the ECD repeats are in dark green and elements at the convex face are in light green. Potential N-linked glycosylation sites of *Ce*LGR as predicted by the NetNGlyc-1.0 server are indicated with blue inverse triangles. N-linked glycosylation sites confirmed in solved structures are highlighted in blue letters: FSHR (PDBid 1XWD) and TSHR (PDBid 3G04). No glycosylation site was reported in the structure of LHCGR (PDBid 7FIH). Cysteine residues are highlighted with yellow background, and disulfide pairs are indicated by orange connections. Residues in P10 are highlighted with red background. Residues subject to proteolytic removal are indicated with grey background. Conserved residues are underscored by asterisks (*).

**Extended Data Fig. 2.**
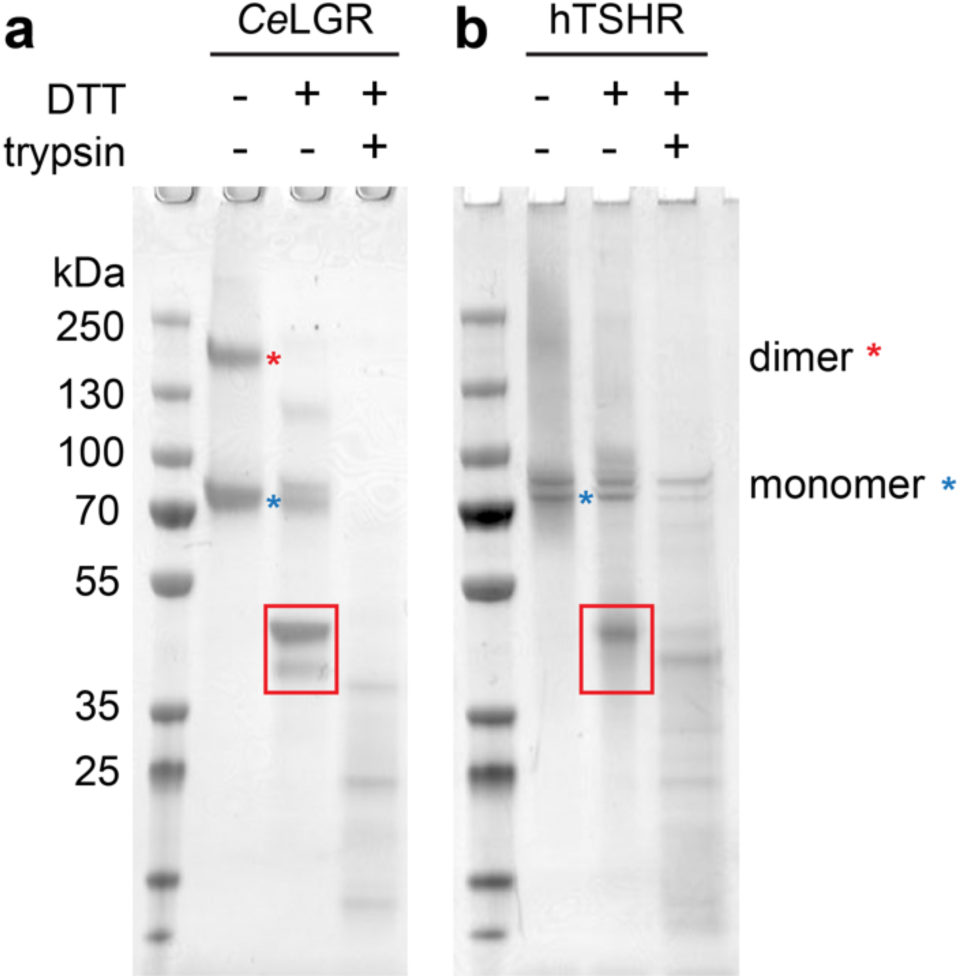
SDS-PAGE analysis of purified receptor proteins. (**a**) **CeLGR**. *Ce*LGR appears as a monomer band and a dimer band in the absence of reducing agent dithiothreitol (DTT). Two new bands corresponding to lower molecular weight appear for *Ce*LGR in the presence of DTT. After limited trypsin digestion, the residual monomer band is almost entirely displaced to smaller fragment bands. **(b) hTSHR.** The putative ‘dimer’ portion of the gel for hTSHR in non-reducing conditions is diffuse. A new band, similar to the one for *Ce*LGR, plus additional diffuse densities appear for hTSHR in DTT. After limited trypsin digestion under the chosen conditions, residual monomer density is much reduced and new densities appear for fragments. The bands excised from the gel for analysis by proteolytic mass spectrometry are boxed in red.

**Extended Data Fig. 3.**
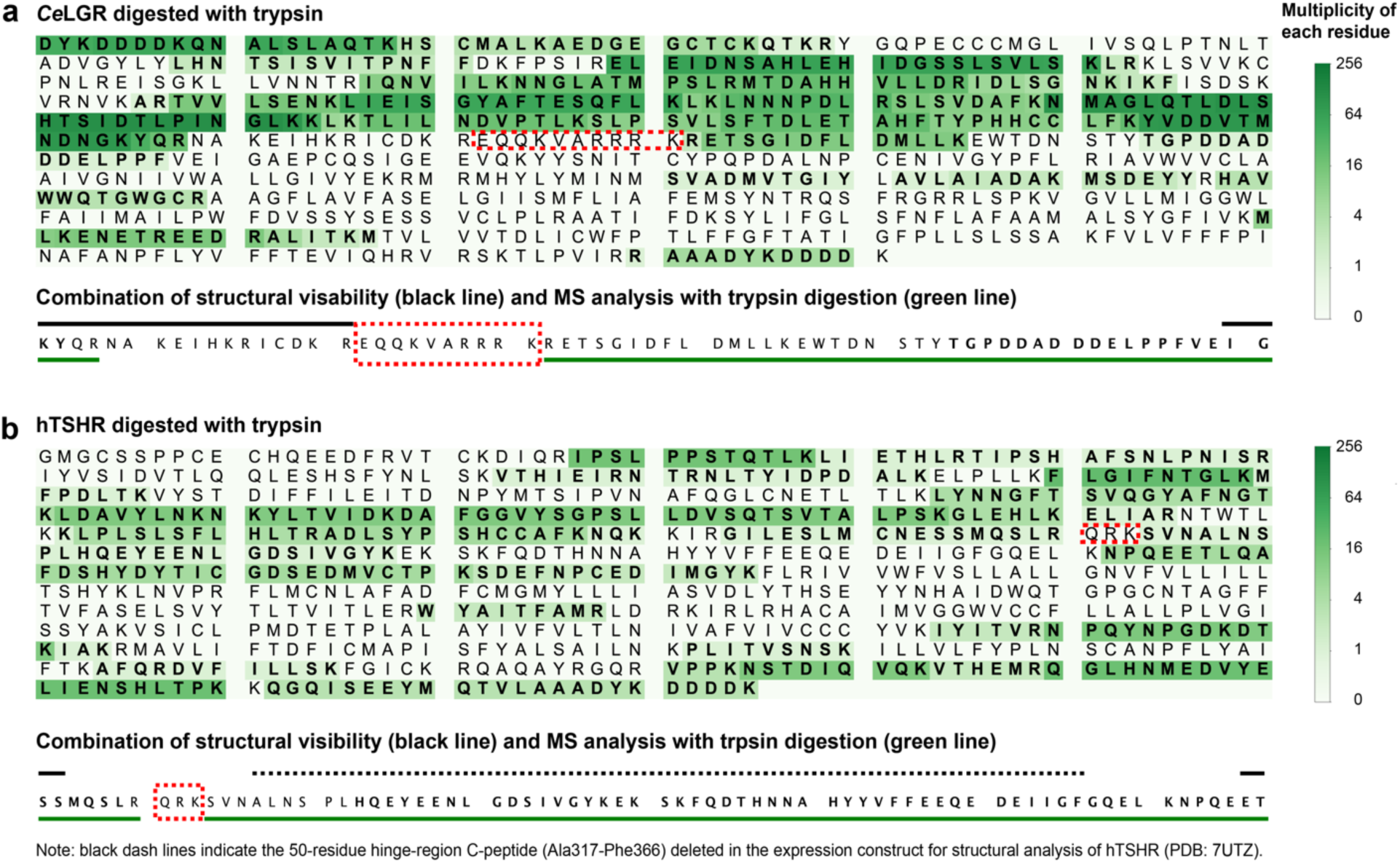
Delimitation of the cleavage site(s) with a combination of proteolytic mass spectrometry and structural coverage. (a) CeLGR. (b) hTSHR. For each receptor and each protease, the multiplicity of tryptic peptides is represented with gradient opacity of color as identified in the color bar at the right. The cleavage efficiencies for chymotrypsin and GluC were lower, giving less coverage from identified peptides; thus, these enzymes did not further delimit the site. The combined results for each receptor are summarized below the peptide maps as colored lines below the delimiting sequence; the span of ordered structure in the respective cryo-EM maps are shown in black above each sequence; and the integrated delimitation of possible cleavage is highlighted in a dashed red box covering the sequence.

**Extended Data Fig. 4.**
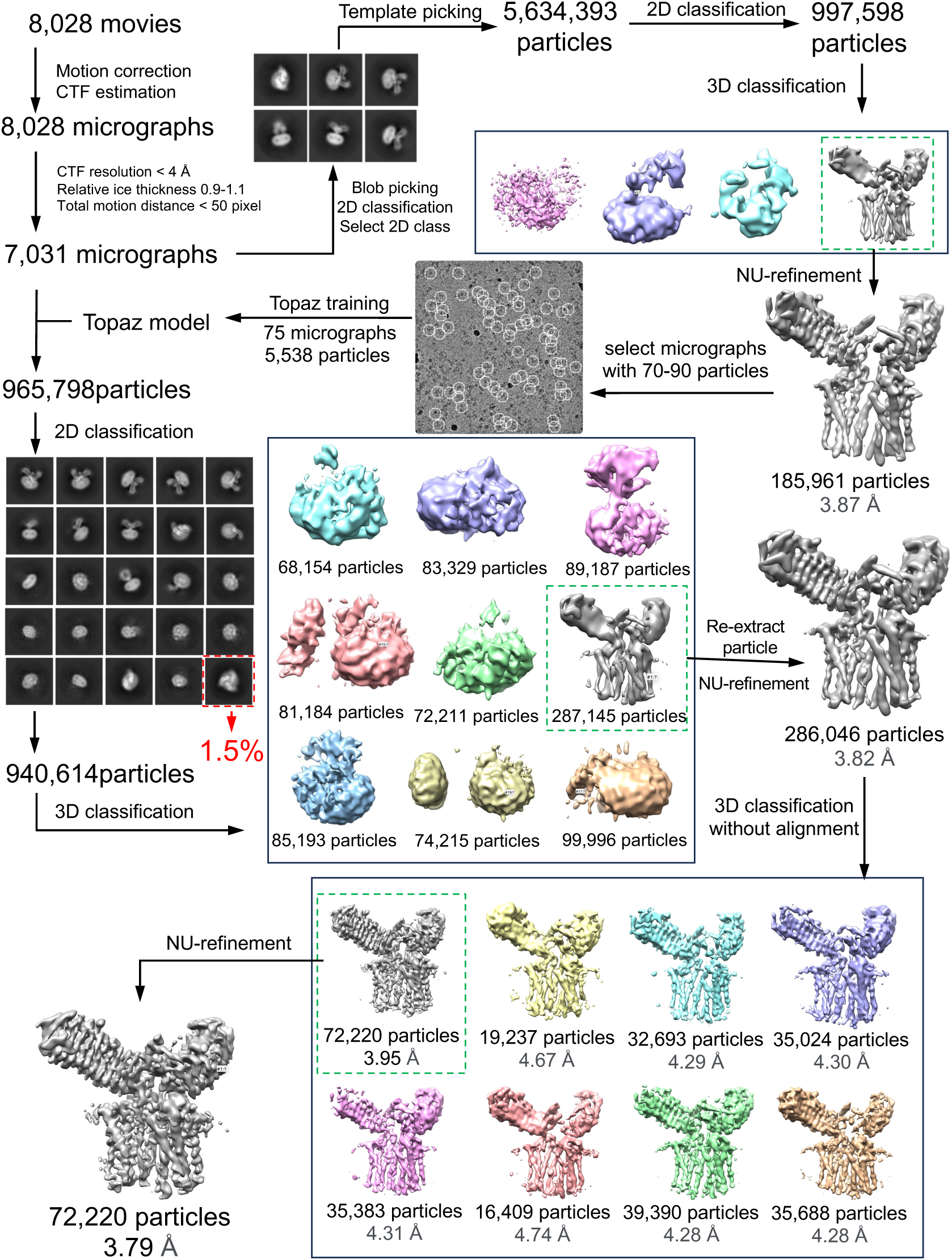
Flowchart outlining cryo-EM image acquisition and processing for the structure of *Ce*LGR homodimer in apo state. The green dashed box displays the chosen high-quality 3D class, while the red dashed box represents a top view of a putative *Ce*LGR trimer. All processing was performed using cryoSPARC (see Methods for details).

**Extended Data Fig 5.**
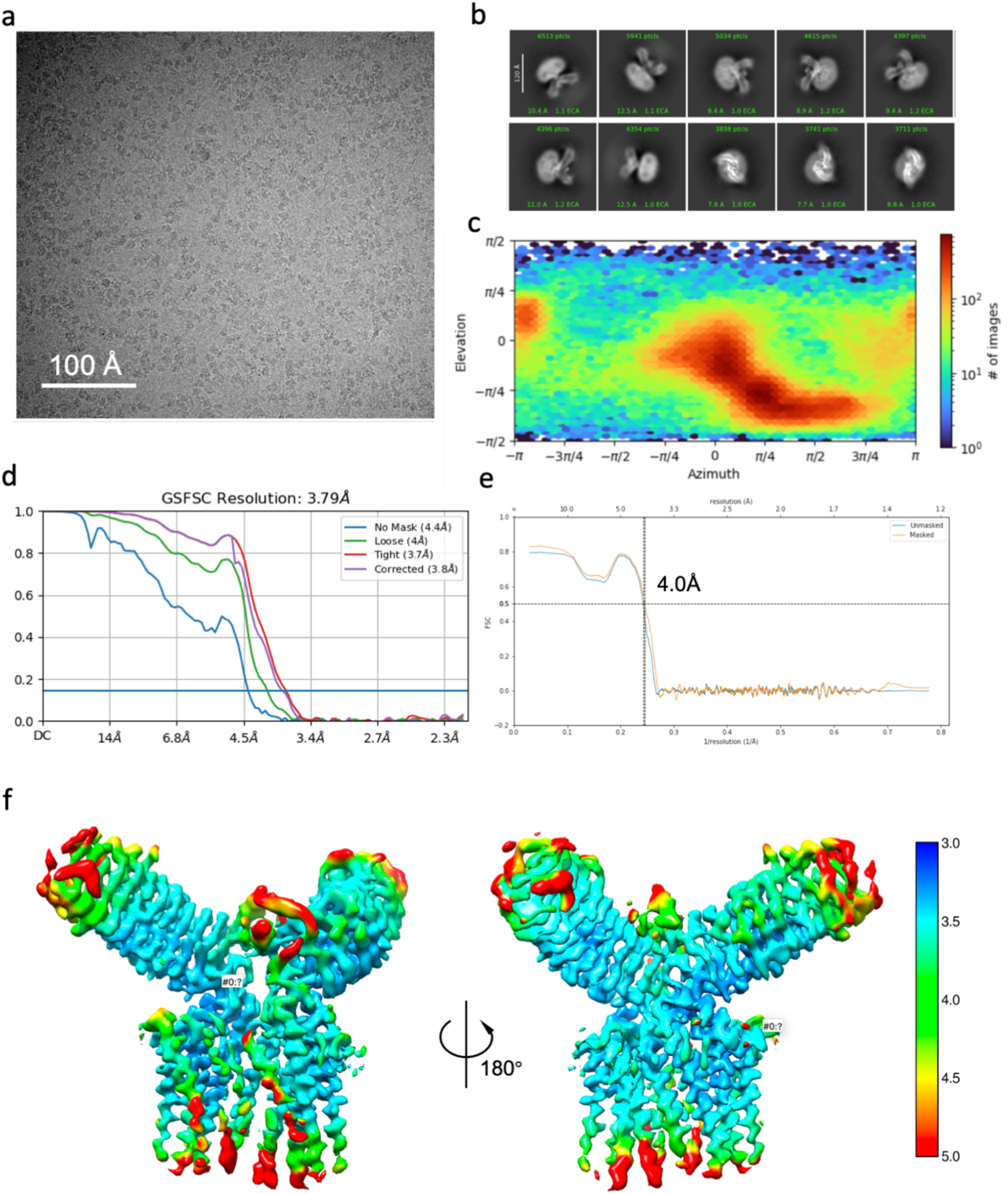
Cryo-EM data analysis for the *Ce*LGR homodimer in apo state. Representative micrograph **(a)** and 2D classes **(b)** for the *Ce*LGR homodimer. **(c)** Heat map of the Euler angle distribution of particle orientations from the final 3D reconstruction in CryoSPARC v4.5.0. **(d)** Fourier shell correlation (FSC) curves between the two independently refined half-maps of final cryo-EM 3D reconstruction. **(e)** Map to model FSC curves. **(f)** Sharpened map of *Ce*LGR homodimer colored by local resolution.

**Extended Data Fig 6.**
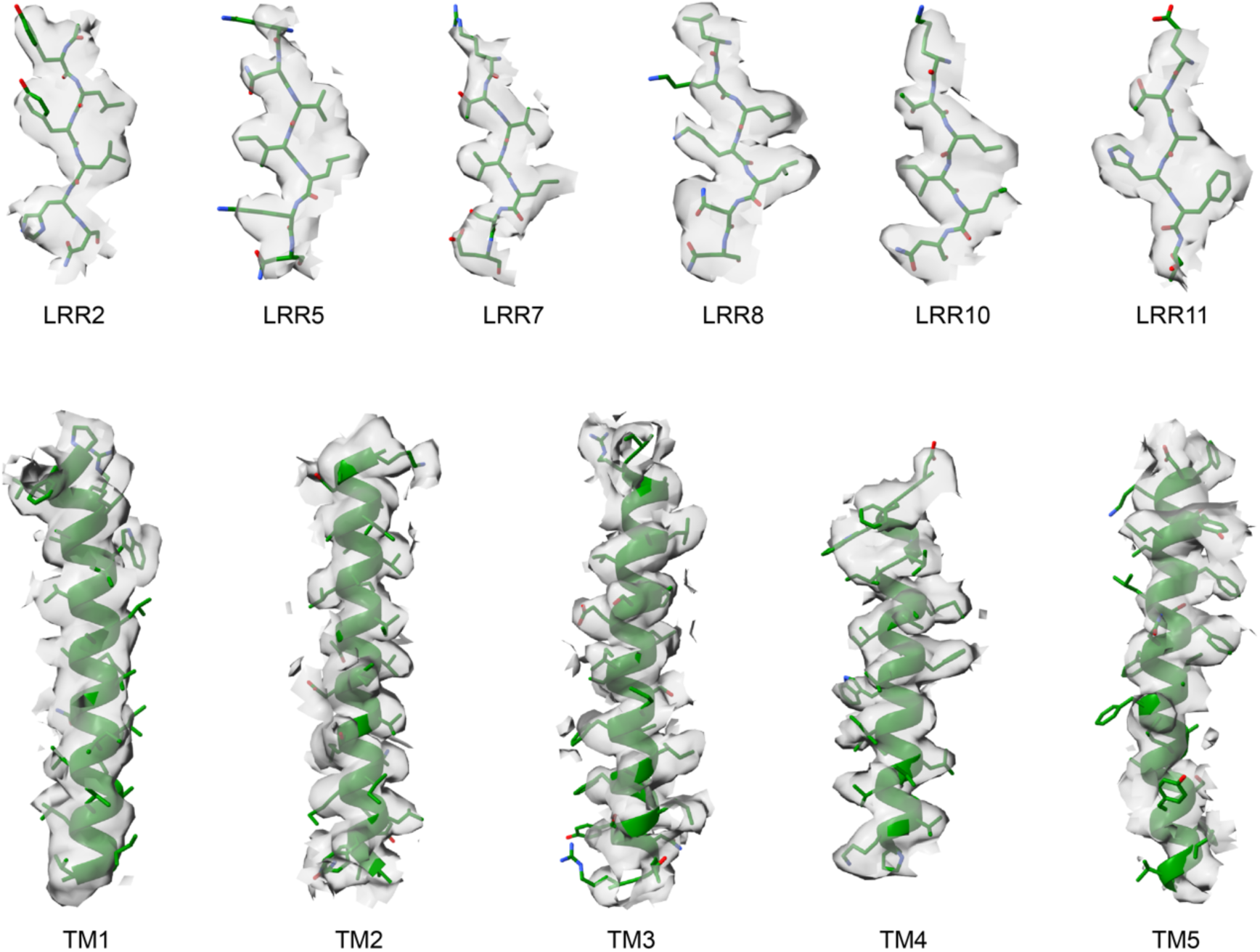
Model-to-map fittings for selected features. Cryo-EM densities (transparent grey surface) are shown with corresponding segments of the atomic model for *Ce*LGR protomer A.

**Extended Data Fig. 7.**
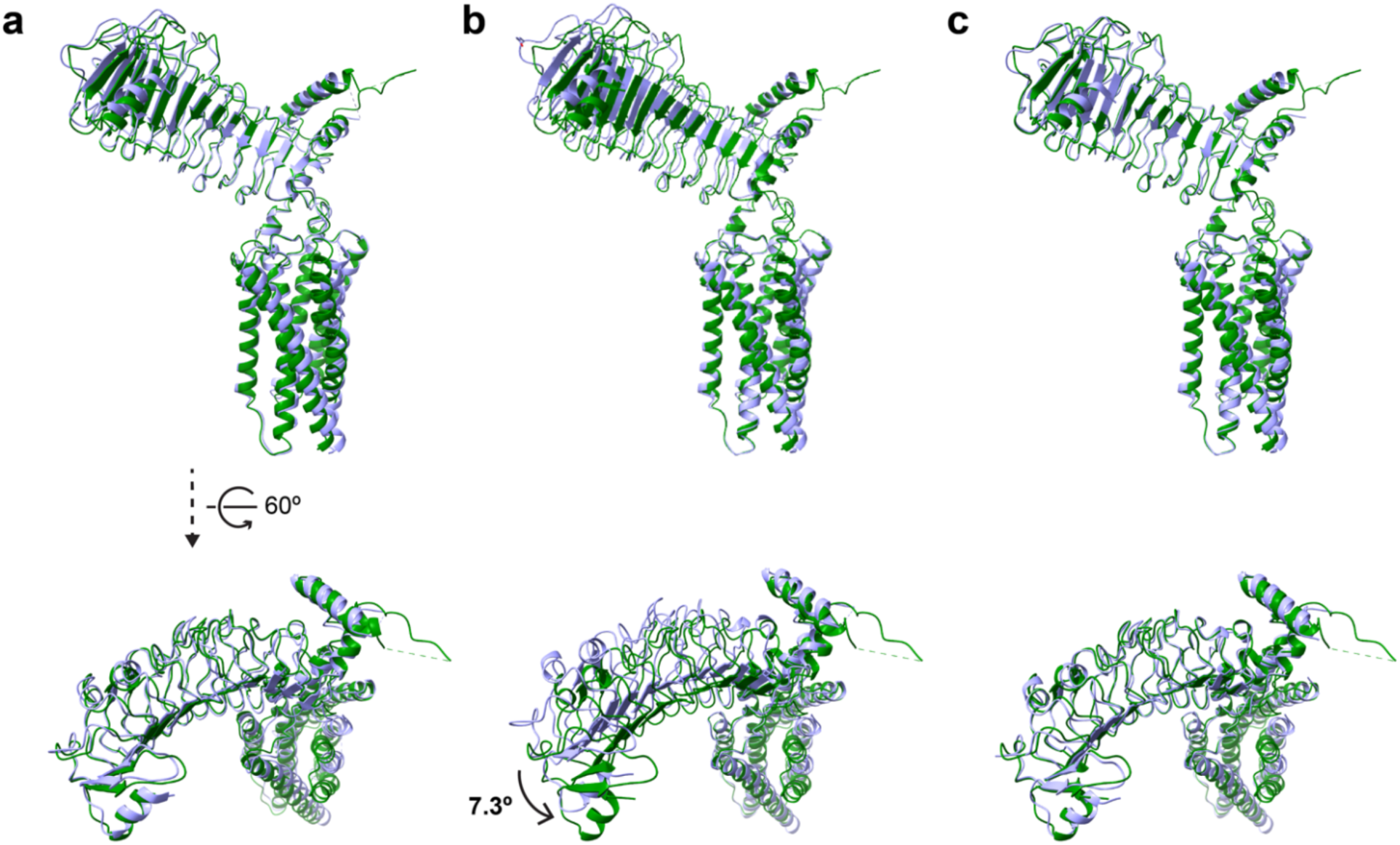
Comparison between protomers A (green) and B (light purple) of the apo-state *Ce*LGR homodimer. **(a)** Overall superimposition of protomer B onto A. **(b)** Superimposition of the TMD of protomer B onto the TMD of protomer A. **(c)** Superimposition of the ECD of protomer B onto the ECD of protomer A after first superimposing the TMDs as in **(b)**.

**Extended Data Fig. 8.**
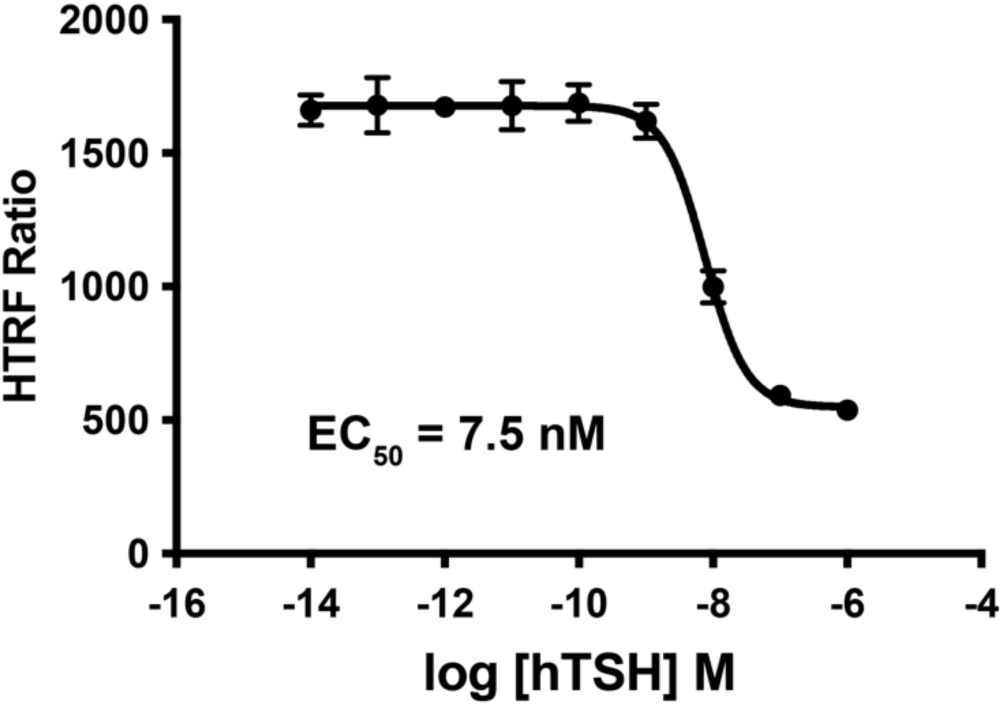
Activation of hTSHR by hTSH. The EC50 of hTSHR activation by hTSH was measured using a cAMP-based competitive assay in HEK293S cells as a control. Data points (mean ± s.d) are plotted for measurements from an experiment of n=3 biological replicates.

**Extended Data Fig. 9.**
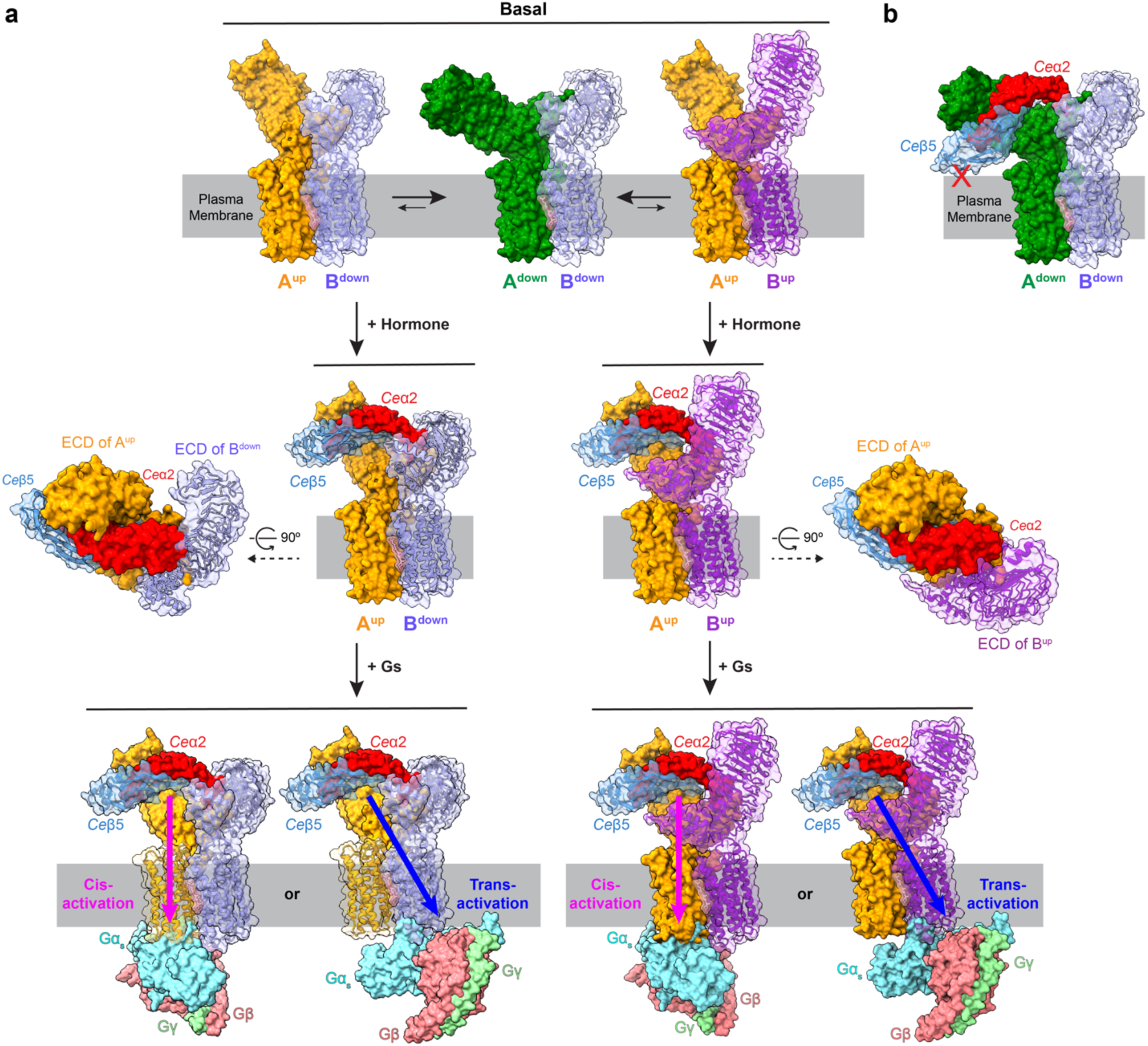
Implications for receptor activation. **(a)** In the basal state, the ECD of unliganded *Ce*LGR can spontaneously transit to the up conformation: either one ECD-up (left) or two ECDs-up (right) seems possible. Binding of *Ce*α2β5 ligand will stabilize the ligand-bound ECD in the ‘up’ conformation. G protein coupling to either protomer A (cis-activation) or protomer B (trans-activation) seems feasible. In each image, protomer A and *Ce*α2 are drawn as opaque surfaces while protomer B and *Ce*β5 are drawn as transparent surfaces exposing ribbon diagrams of the polypeptide backbone. **(b)** Binding of *Ce*α2β5 onto a *Ce*LGR dimer in the ‘down’ conformation might cause collision between the *Ce*β5 subunit and cell membrane.

**Extended Data Table 1.**
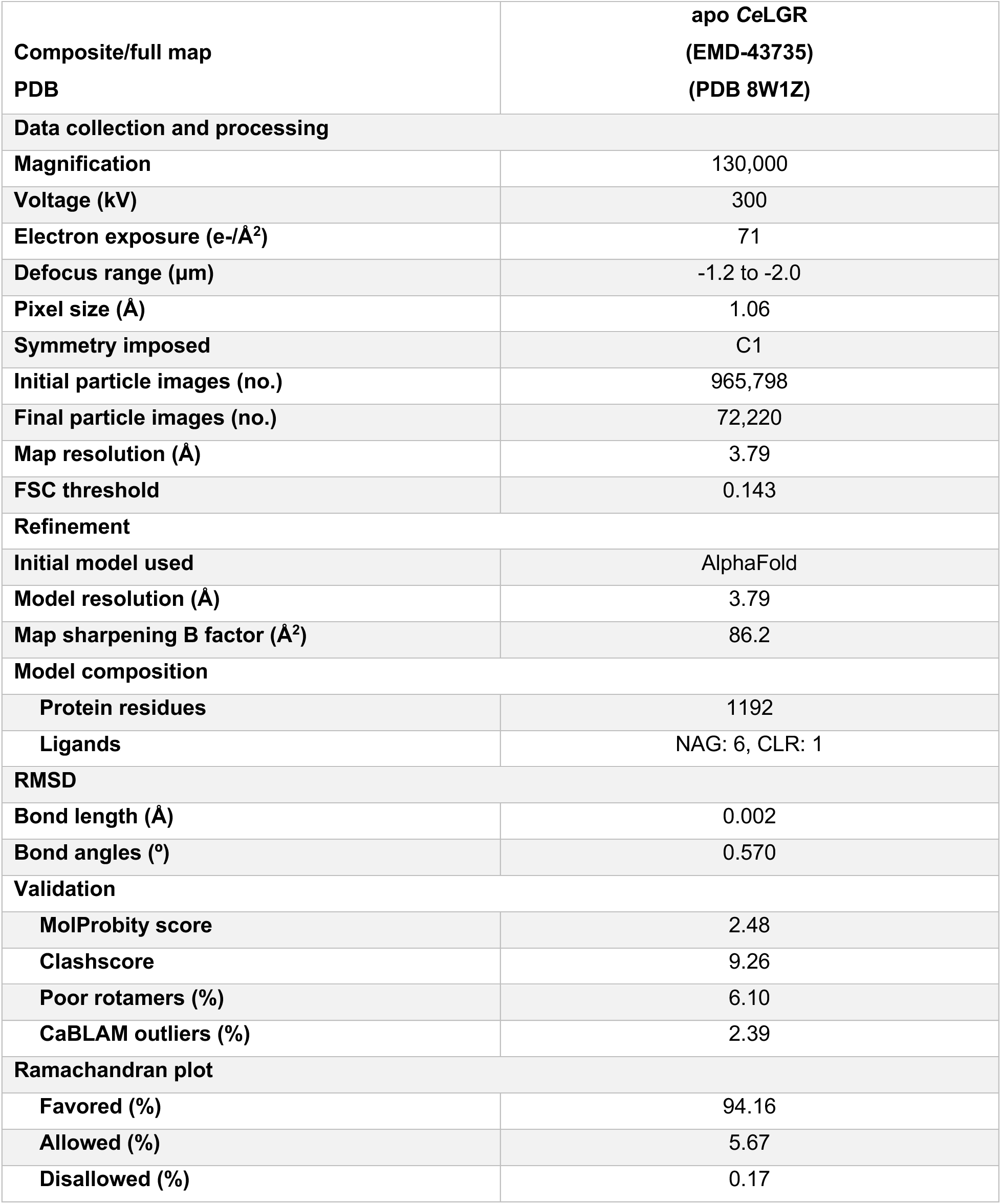
Cryo-EM data collection, refinement, and validation statistics

**Supplementary Video 1. Top view of the global conformational variability analysis of *Ce*LGR homodimer in apo state.** The angle between the ECDs of protomer A (left) and protomer B (right) changes continuously. Simultaneously, the TMDs contract and extend like a ‘heart’.

**Supplementary Video 2. Side view of the global conformational variability analysis of *Ce*LGR homodimer in apo state.** Protomer A is on the left and protomer B is on the right.

